# CaptureBody – an anti-CD45 x anti-IgG bispecific antibody enables accurate unmixing for spectral flow cytometry

**DOI:** 10.64898/2026.02.13.704926

**Authors:** Alexander E. Zambidis, Latha Kallur Siddaramaiah, Andrew J. Konecny, Matthew Gray, Martin Prlic

**Affiliations:** Vaccine and Infectious Disease Division, Fred Hutchinson Cancer Center, Seattle, WA, USA; Department of Immunology, University of Washington, Seattle, WA, USA; Department of Global Health, University of Washington, Seattle, WA, USA

## Abstract

Accurate spectral unmixing is a critical step for flow cytometry data analysis and requires a single stain control for every fluorescent parameter used in an experiment. Currently, compensation particles are often used for making single stain controls when a target protein is of low abundance or a cell type is of low frequency. However, compensation particles introduce incongruencies in emission spectra compared to cells resulting in spectral unmixing or compensation errors. To enable the use of cells regardless of the abundance of target proteins or immune cell type, we generated a bispecific antibody that links a human anti-CD45 and mouse anti-IgG variable region. We refer to this new bispecific tool as CaptureBody (CB) and highlight the benefits of its final nanobody-based design. We provide all sequences and methods necessary for the in-house expression of a CaptureBody to disseminate their use for spectral flow cytometry experiments.

## Introduction

Recent advances in spectral flow cytometer instrument design^1^, development of new fluorescent dyes^2^, and improved methods for spectral unmixing^3,4^ have opened the possibility of concomitantly increasing data quality and the number of parameters that can be measured. These advancements allow for highly complex staining panels, currently up to 50 fluorochromes, that provide in-depth analysis of immune cell subsets and their activation status.^5,6^ Importantly, such high parameter panels depend on the implementation of accurate single stain controls to prevent modeling error within the final unmixed data.^7^

Briefly, the hallmark feature of spectral flow cytometry is the capture of the full emission spectrum of a fluorochrome across multiple lasers in contrast to only capturing specific, narrow wavelengths (as is done in conventional flow cytometry). Single stain spectra are the basis for subsequently “deconstructing” the complex spectral emission of a cell stained with possibly dozens of antibodies to accurately extrapolate signal identity and strength of each fluorochrome.^8,9^ Thus, accurate single stain controls are critically important to capture the full emission spectrum for each fluorochrome.

For human studies, healthy donor PBMCs are often used to create single stain controls because they are an accessible and abundant source of cells. However, healthy donor PBMCs can differ from samples of interest (such as human cohorts with immune alterations, tissue biopsies, etc.) and cannot be used when the target protein is expressed at a lower antigen density on PBMCs than any of the samples, or when the cell type that expresses the target protein is absent/rare in PBMCs. In these situations, compensation/unmixing particles are used to generate single stain controls as they provide a positive population with a bright and uniform signal.

However, there are drawbacks to using compensation/unmixing particles. One challenge is that certain fluorochromes produce spillover coefficients or normalized spectra that differ when bound to a compensation/unmixing particle compared to a cell.^5,10,11^ These differences result in inaccurate fluorescence modeling for compensation or spectral unmixing. The consequences can range from a loss of signal resolution to the introduction of errors within the final data set.^5,10,11^

Here, we present a novel tool to bypass the need for compensation/unmixing particles and enable the use of PBMCs for all single stain controls. We accomplished this by generating a bispecific antibody that can bridge all human immune cells to commercially available fluorochrome-conjugated antibodies. In research and clinical applications, bispecific antibodies have grown popularity due to their highly customizable size, structure, antigen specificity, and valency.^12^ Bispecific antibodies have been developed to facilitate receptor activation or inhibition, mimic co-factors, sequester soluble molecules such as growth factors and cytokines, allow for piggybacking into tissue or cellular compartments otherwise inaccessible, and heighten selectivity for ADCC and antibody drug conjugates.^13^ Additionally, bispecific antibodies have been designed to bridge effector immune cells with malignant tumors to exert targeted cytotoxic effects typically inhibited by the tumor microenvironment.^14^ This ability to bridge two surfaces was an attractive property for why we selected a bispecific as the bases for our novel reagent.

We initially developed and tested three different formats of bispecific reagents to remove the current limitations of cell-based single stain controls for flow cytometry. Since the bispecific antibody binds to immune cells and the provided single stain antibody, we refer to this these reagents as a CaptureBody. We engineered and tested three CaptureBody variants using a combination of previously published nanobody and scFv sequences to create novel Fc-less bispecific reagents with a small size profile. We suggest that our CaptureBody-3 variant that binds to CD45 and the kappa chain of mouse IgG may be best suited for most applications.

We provide all CaptureBody sequences and expression methods to facilitate common implementation of this new tool. Importantly, our work provides a new avenue to capture accurate emission spectra for single stain controls which until this point has been a bottleneck for spectral unmixing and will also be beneficial for complex flow cytometry panels on conventional cytometers.

## Results

### Initial CaptureBody design considerations, design approach and generation

We considered that the bispecific reagent must uniformly stain cells and bind to a cell surface protein that is highly expressed. We were also concerned that a large bispecific would introduce steric hinderance and reduce staining. Therefore, we considered two distinct approaches to designing a bispecific reagent with a small size profile. The first was to isolate the variable heavy and light chain fragments from an antibody, and connect them with a short peptide linker, called a single-chain variable fragment (scFv). This approach retains original antigen specificity and allows for modular design in therapeutics, diagnostic tools, or laboratory reagents.^15^ The second was to isolate the variable region from camelid single-domain antibodies, called nanobodies or V_HH_.^16,17^ In emerging technologies, nanobodies demonstrate many advantages over scFv antibody fragments due to their high solubility, low immunogenicity, and ease of modular design without the need for light chain association.^18,19^ In addition to retaining antigen specificity, the relative small size of nanobodies (∼15kDa) allows for access to non-planar surfaces typically inaccessible to larger scFvs (∼30kDa) and full human antibodies (∼150kDa).^20^ We decided to implement and compare both approaches throughout the development of each CaptureBody in addition to removing the Fc region to further decrease size and remove any potential aggregation by self-reactivity.

To design the CaptureBody reagents, we utilized variable region sequences from publicly available scientific and patent literature. Pleiner et al. first developed a collection of anti-mouse and anti-rabbit IgG nanobodies with validated epitopes and cross-species specificity. These nanobodies were generated by immunizing alpacas and selected by a phage display library.^21^ They were originally designed as secondary labeling reagents for western blot, confocal, and super-resolution microscopy that outperformed conventional secondary antibody labeling. From this toolbox, we selected clone TP1107 which is specific for mouse IgG1 Fc with species cross-reactivity to rat and human for the antibody capture domain used in CaptureBody-1 (CB-1) (**Figure 1A**). For the cell binding domain in CB-1, we created a scFv derived from the variable regions of apamistamab, described in patent US20220175951A1^22^ (**Figure 1A**). This human anti-CD45 antibody has been described as effective to target hematopoietic stem cells before stem cell transplantation when conjugated with the radioconjugate ^131^I.^22,23,24^ We ensured correct variable heavy and light chain association in our human anti-CD45 scFv by connecting the heavy and light variable fragments via a flexible glycine-serine linker (G4S)_4_ (**Figure 1A**). To join the two binding domains, we used a GCN4 spacer to allow for oligomerization with two 6xHis tag inserts to allow for nickel chromatography purification (**Figure 1A**).

**Figure 1:**
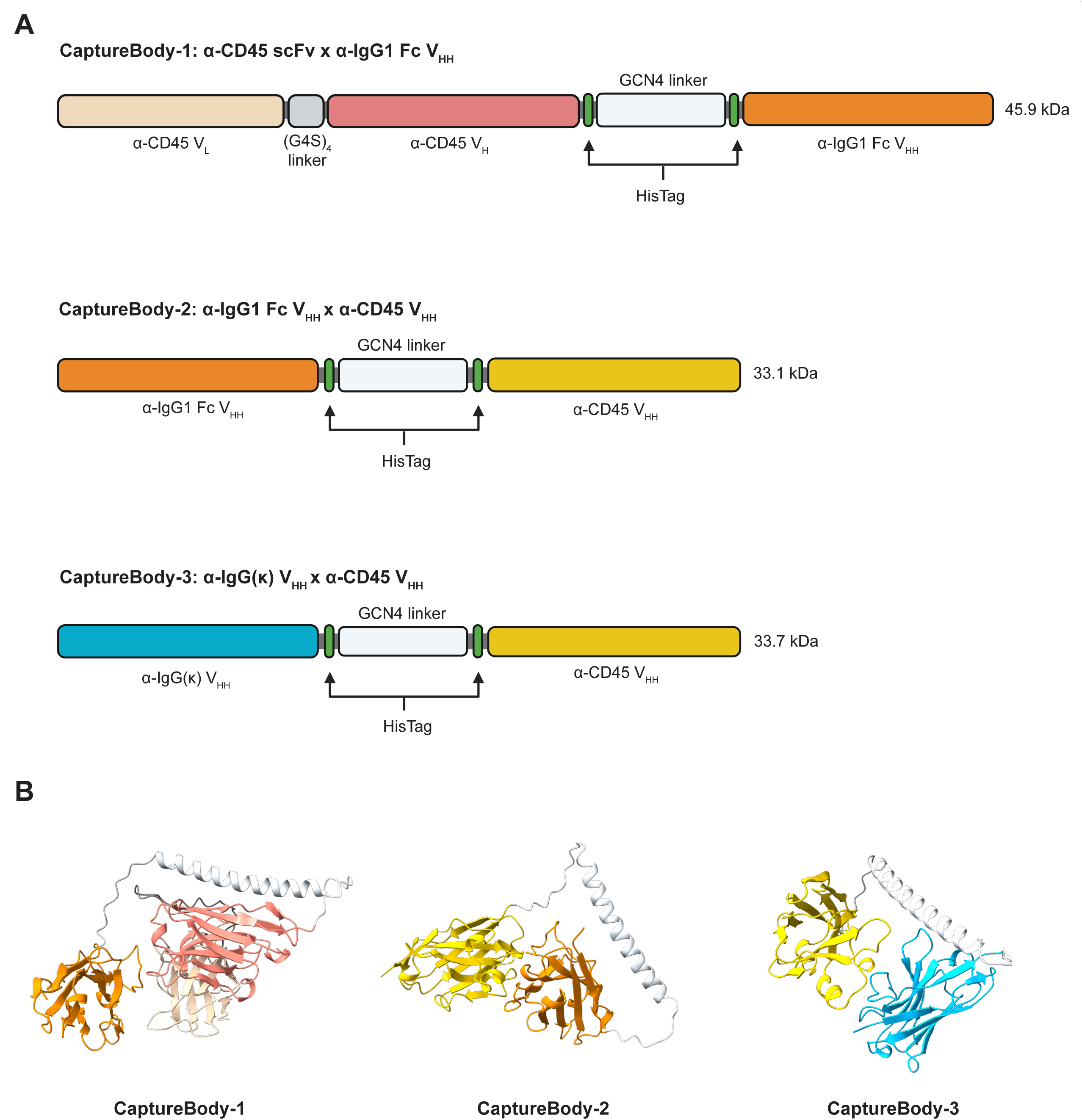
CaptureBody design and AlphaFold3 modeling. (A) Schematic diagram of the amino acid sequence of each CaptureBody with monomer molecular weight. Specific regions are delineated by color: anti-CD45 scFv variable light chain (beige), anti-CD45 scFv variable heavy chain (red), scFv linker (gray), HisTag (green), GCN4 linker (silver), anti-IgG1 Fc nanobody (orange), anti-CD45 nanobody (yellow), anti-IgG(κ) nanobody (blue). (B) AlphaFold3 structure prediction of each monomeric CaptureBody. CB-1 (left), CB-2 (middle), and CB-3 (right)

For the creation of our second CaptureBody (CB-2), we wanted to find an alternative for using an scFv for the cell binding domain. We identified clone 2H5, a human anti-CD45 nanobody derived from patent US20230357424A1^25,26^, as a candidate for our CaptureBody design. Clone 2H5 was derived by immunizing alpacas with human CD45 and selected via phage display library. 2H5 was epitope mapped using surface plasmon resonance and found to specifically target the N-terminal D2 extracellular domain of CD45. Lupardus & Rokkam discovered several nanobodies with specificity for all distinct sites across the four CD45 fibronectin type III-like extracellular domains to provide new tools for studying the role of CD45 in T cell function, as nanobodies targeting distinct regions of CD45 may prove useful in downstream experimentation.^27,28^ Here, we kept TP1107 for the antibody capture domain of CB-2, however, it is important to note that for the CB-1 design, TP1107 is on the C-terminus while for the CB-2 design TP1107 is on the N-terminus. Additionally, the use of nanobodies for both binding domains in CB-2 allowed for a ∼12kDa reduction in CaptureBody size.

Finally, we considered that not all mouse fluorescent-conjugated antibodies are of IgG1 isotype, however, the vast majority contain kappa light chains. Thus, we designed CB-3, which included Clone 2H5 for the cell binding domain as in CB-2, but instead used the kappa chain-specific clone TP1170 from Pleiner et al. for the antibody capture domain (**Figure 1A**). Therefore, using TP1170 would allow for broader applicability of CB-3 in flow cytometry. Like in CB-2, the antibody capture domain TP1170 of CB-3 was placed on the N-terminus (**Figure 1A**).

As a proof of principle for design, we first used AlphaFold3 (AF3) to model all CaptureBodies as monomers *in silico* to explore whether the cell binding and antibody capture domains showed disorganization when linked together. Based on the promising AF3 predictions (**Figure 1B**), we proceeded with the production of each CaptureBody. Next, we characterized their oligomerization and binding properties to their respective antigen.

### CaptureBody characterization and optimization

To assess individual antigen binding, we performed biolayer interferometry (BLI) against human CD45 and mouse IgG (**Figure 2A**). While both tandem nanobody CaptureBodies CB-2 and CB-3 exhibited lower total binding, they reached saturation faster and remained bound longer to their respective antigen through their dissociation phase when compared to CB-1 (**Figure 2A**). As previously mentioned, CB-1 and CB-2 have the same antibody capture domain, however, CB-1’s is located on the C-terminus while CB-2’s is located on the N-terminus. It is plausible that this difference in orientation of the antibody capture domain led to the observed difference in IgG binding. Each CaptureBody was then purified by nickel affinity chromatography, filter sterilized, and run over size exclusion chromatography (SEC) to assess oligomerization state. Initially, the GCN4 domain linker in CB-1 and CB-2 produced distinct dimer and trimer populations (**Figure 2B**). To reflect the natural homodimer formation found in IgGs, we sought to drive the preferential production of dimer over trimer. Computationally outlined by Volkmer and Hochreiter et al., specific amino acid substitutions in GCN4 can influence dimer or trimer oligomerization.^29^ Before GCN4 linker optimization, CB-1 expressed primarily as a trimer with a dimer subpopulation while CB-2 expressed primarily as dimer with a trimer subpopulation (**Figure 2B**). After inserting a modified GCN4 linker via DNA mutagenesis with the amino acid substitutions I8V, I15N, I22K, K26E, and I29V (**Table S1**), CB-1 expressed with a larger dimer population while CB-2 exclusively expressed as a homogenous dimer (**Figure 2B**). CB-3 was only produced with the modified GCN4 linker and exhibited a sole dimer peak (**Figure 2C**). We next ran SDS-PAGE under reducing conditions to confirm monomeric unit size, with each bispecific matching its designed monomer molecular weight (**Figure 2D**). All SEC purified dimer fractions were collected and used for subsequent flow cytometry evaluation. Our characterization results demonstrated that individual antigen binding occurred in the bispecific format across all CaptureBodies.

**Figure 2:**
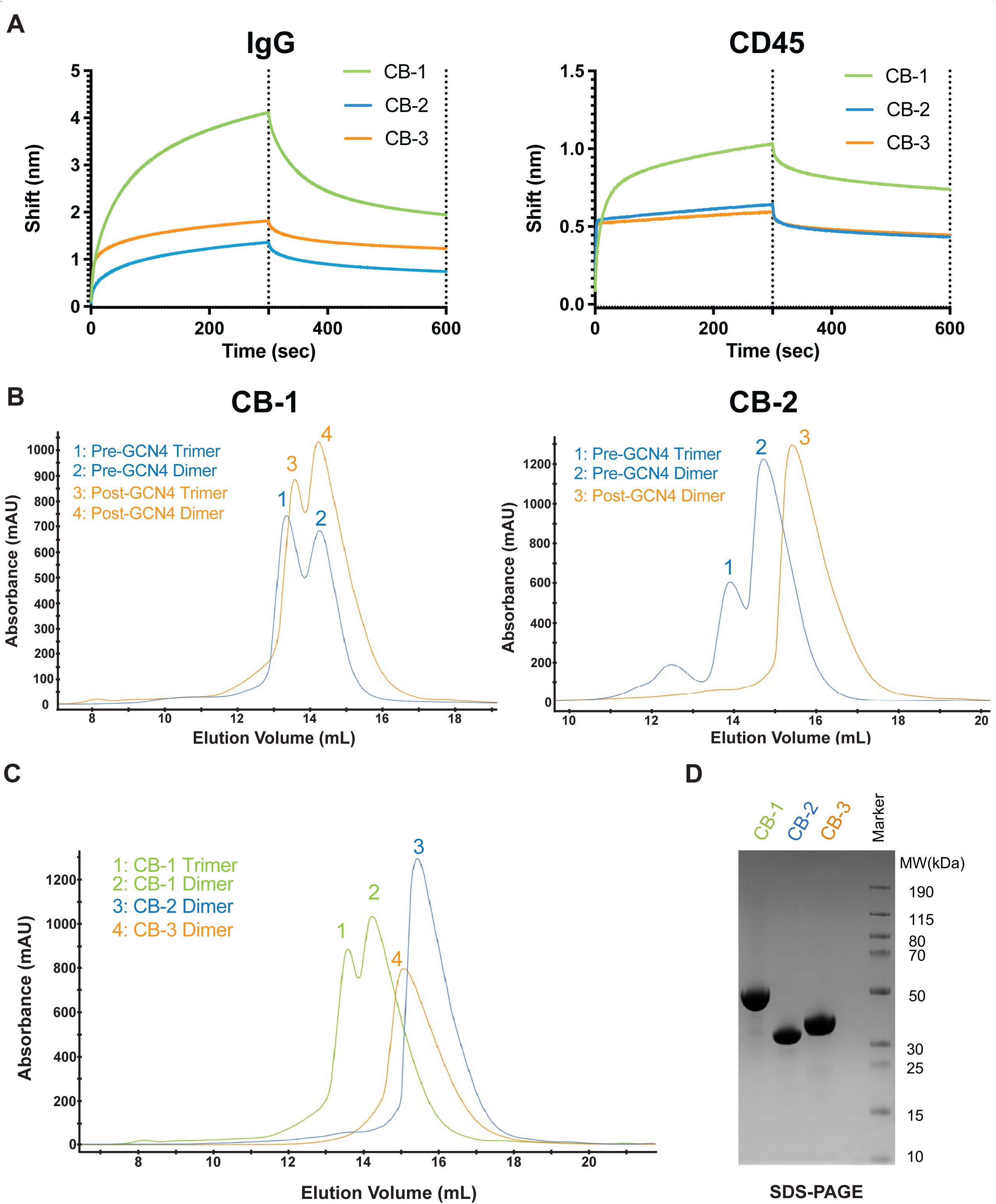
CaptureBody characterization and optimization. (**A**) Bio-layer interferometry data assessing binding to IgG and CD45. Fc probes were pre-loaded with either mouse serum (40μg/mL) or CD45-Fc (2uM) then dipped into 2uM of each CaptureBody in conditioned media following collection from HEK293 cells. On-and off-rate recorded through their respective association phase (0-300 seconds) and dissociation phase (300.01-600 seconds). (**B**) Size exclusion chromatography peaks for CB-1 and CB-2 pre- (blue) and post-modification (orange) of GCN4 linkers. (**C**) Size exclusion chromatography peaks of CB-1 (green), CB-2 (blue), and CB-3 (orange), post-GCN optimization. 5mg of each CaptureBody was run in Tris-Buffered Saline (TBS) and dimer fractions were collected for flow cytometry binding. (**D**) SDS-PAGE under reducing conditions to quality control monomeric size of each CaptureBody

### All CaptureBody formats exhibit functional bridging

To determine our CaptureBodies utility in generating single stain controls, we next evaluated their efficacy as a bridging molecule between human PBMCs and fluorochrome-conjugated antibodies. To do this, we stained PBMCs with each CaptureBody at saturating conditions followed by a mouse PE isotype control (isotype IgG1), which by itself would not specifically bind to human cells. Separately, we also directly stained PBMCs at saturating conditions with an anti-CD45 PE antibody as a positive control. PE-conjugated antibodies tend to have a one-to-one fluorochrome to antibody chemistry^30,31^ which we specifically chose for this comparison to limit variation that occurs in labeling antibodies with a fluorescent probe. By using the same mg/mL quantity of mouse PE isotype control and anti-CD45 PE antibody, we were able to directly interrogate whether the quantity of PE conjugated antibodies on the cell surface differed between the commercial anti-CD45 PE antibody and our CaptureBody. All PBMCs stained with our CaptureBody variants exhibited a unimodal peak with a high median fluorescence intensity (MFI) that was comparable to the commercial human CD45 PE antibody (**Figure 3**). Compared to the MFI produced by staining PBMCs with the commercial anti-CD45 PE antibody, CB-1 retained 72.4%, CB-2 retained 82.1%, and CB-3 retained 79.1% of the MFI (**Table S2**). We also performed a serial dilution of each CaptureBody with the mouse PE isotype control (**Figure S1**). These data demonstrate that each CaptureBody can bridge fluorochrome-conjugated antibodies to cell surface CD45. Moreover, CaptureBodies stained cells in a uniform and high antigen density manner, similar to direct staining of CD45. Of note, this is a key requirement for its intended use as a single stain control.

**Figure 3:**
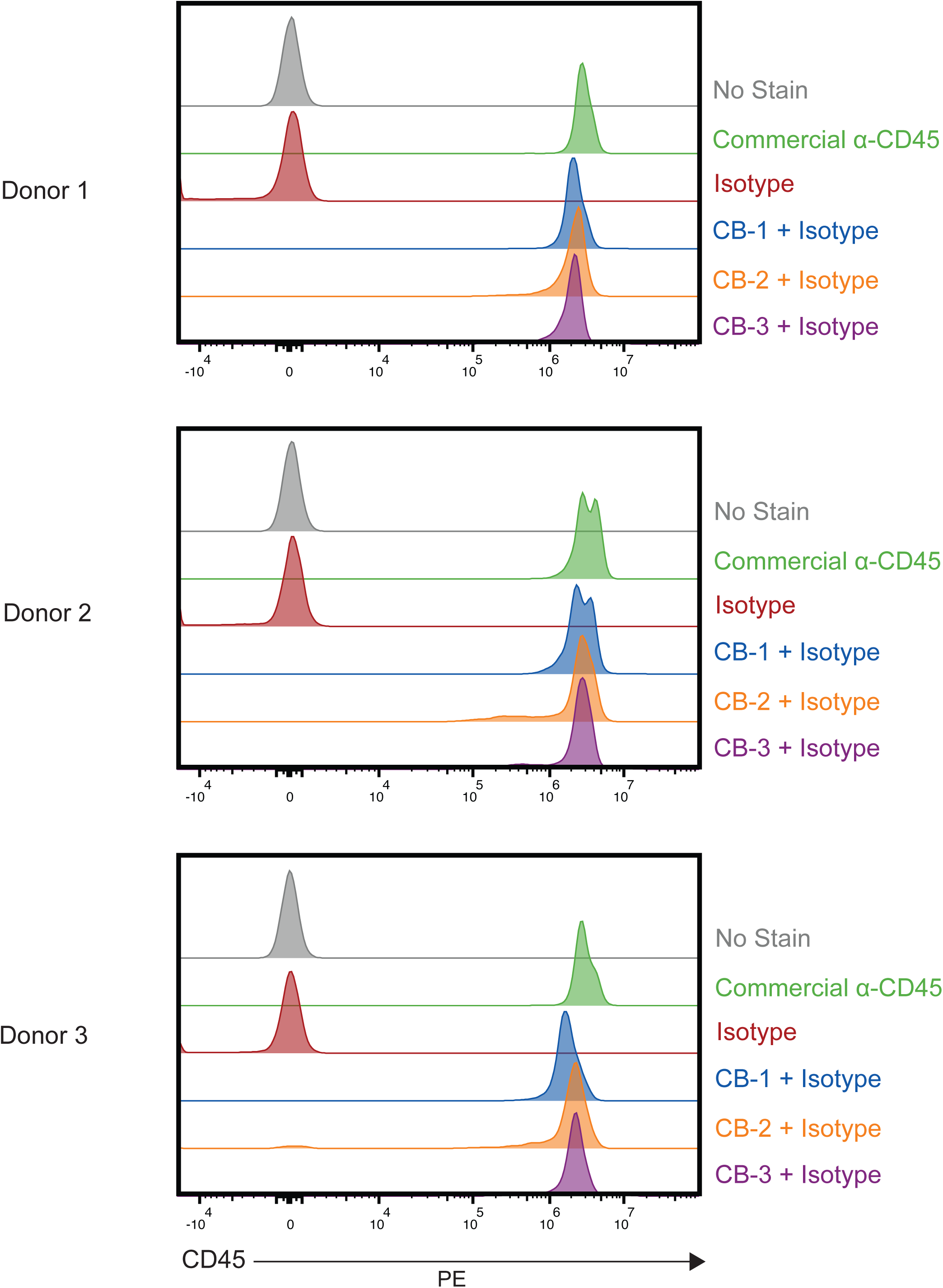
All CaptureBody formats exhibit functional binding. Staining of human PBMCs for fluorescent PE in the YG1 channel on a BD FACSDiscover™ S8 Cell Sorter. Cells were either stained with nothing (gray), a commercial CD45 human antibody (green), a PE mouse control isotype antibody (red), CB-1 and the PE isotype control antibody (blue), CB-2 and the PE isotype control antibody (orange), or CB-3 and the PE isotype control antibody (purple) across three donors

### CaptureBodies recapitulate normalized fluorescence of directly stained cells

To assess the utility of CaptureBodies in generating single stain controls, we compared directly stained PBMCs to CaptureBody-stained PBMCs using anti-CD4 fluorochrome-conjugated antibodies. Importantly, to assess CaptureBody-mediated antibody binding in the absence of direct binding of anti-CD4 antibody to CD4 expressing cells, we depleted CD4 positive cells from PBMCs by magnetic activated cell sorting. We also included compensation/unmixing particles to directly assess the benefit of using a CaptureBody in context of fluorochromes that are known to cause problems when used with compensation/unmixing particles (BUV661^10^, BUV737^5^, BV711^5^, PerCP-Cy5.5^11^) as well as a fluorochrome that is reported to have no issues (PE-Cy5^5,11^). Next, we stained these CD4 depleted PBMCs with each of our CaptureBodies followed by staining with an anti-CD4 antibody (either BUV661, BUV737, BV711, PerCP-Cy5.5, or PE-Cy5). The same anti-CD4 antibody was also used to stain donor matched non-depleted PBMCs (that contain CD4 T cells), and compensation/unmixing particles (**Figure 4A**). All samples were then acquired on a BD FACSDiscover S8 Cell Sorter, a high-end spectral flow cytometer equipped with 5 lasers (349 nm, 405 nm, 488 nm, 561 nm, and 637nm) and 78 APDs for detection of fluorescence across the full visible spectrum of light (375 nm to 845 nm). For generating the normalized spectra values, the same gate for the positive population was used for all groups within a given fluorescent probe to control for differences in normalized spectra due to brightness (**Table S3)**. As previously reported, BUV661, BUV737, BV711, and PerCP-Cy5.5 had altered normalized spectra when bound to compensation/unmixing particles, while PE-Cy5 performed well (**Figure 4B**). Normalized spectra differences across all channel detectors can be further examined via cosine similarity.^32,33^ Here, each spectrum is represented as a 78-dimensional vector, corresponding to the recorded emission across each of the 78 detectors on the FACSDiscover S8 cytometer. In flow cytometry applications, the resulting cosine between two emission vectors ranges from 0 (no spectral overlap) to 1 (complete spectral overlap). Using cosine similarity, we compared every alternative single stain control (CB-1, CB-2, CB-3, compensation/unmixing particles) to directly stained PBMCs. All CaptureBody-derived controls had a cosine similarity value greater than 0.999 when compared to directly stained PBMCs, demonstrating high similarity when considering all detectors. The lowest cosine similarity value for each CaptureBody control was for PerCP-Cy5.5; CB-1 (0.99915), CB-2 (0.99990), and CB-3 (0.99984). All controls made using compensation/unmixing particles had a cosine similarity value below 0.999 except for PE-Cy5 (0.99986). The lowest cosine similarity value using compensation/unmixing particles was for BV711 (0.99125) (**Figure 4C**). By each individual detector for the BV711 control, compensation/unmixing particles greatest percent difference was for detector R3 (700 nm) with a difference of 21.34% when compared to directly stained PBMCs (**Figure 4D**). For the same detector, CB-3 BV711’s only differed by 0.25% compared to directly stained PBMCs (**Figure 4D**). Of note, the greatest percent difference for CB-3 BV711 was for detector UV4 (440 nm) at 0.32 % (**Figure 4D**). Moreover, the greatest percent difference for a detector observed for CB-3 across all controls was for the PerCP-Cy5.5 control for detector R3 (700 nm) with a delta of 2.24% compared to directly stained PBMCs (**Figure S2-3**). Together, these data illustrate that CaptureBody single stain controls recapitulate the normalized spectra of directly stained single stain cell controls with high precision, even with fluorochromes that have been shown to be problematic when bound to compensation/unmixing particles.

**Figure 4:**
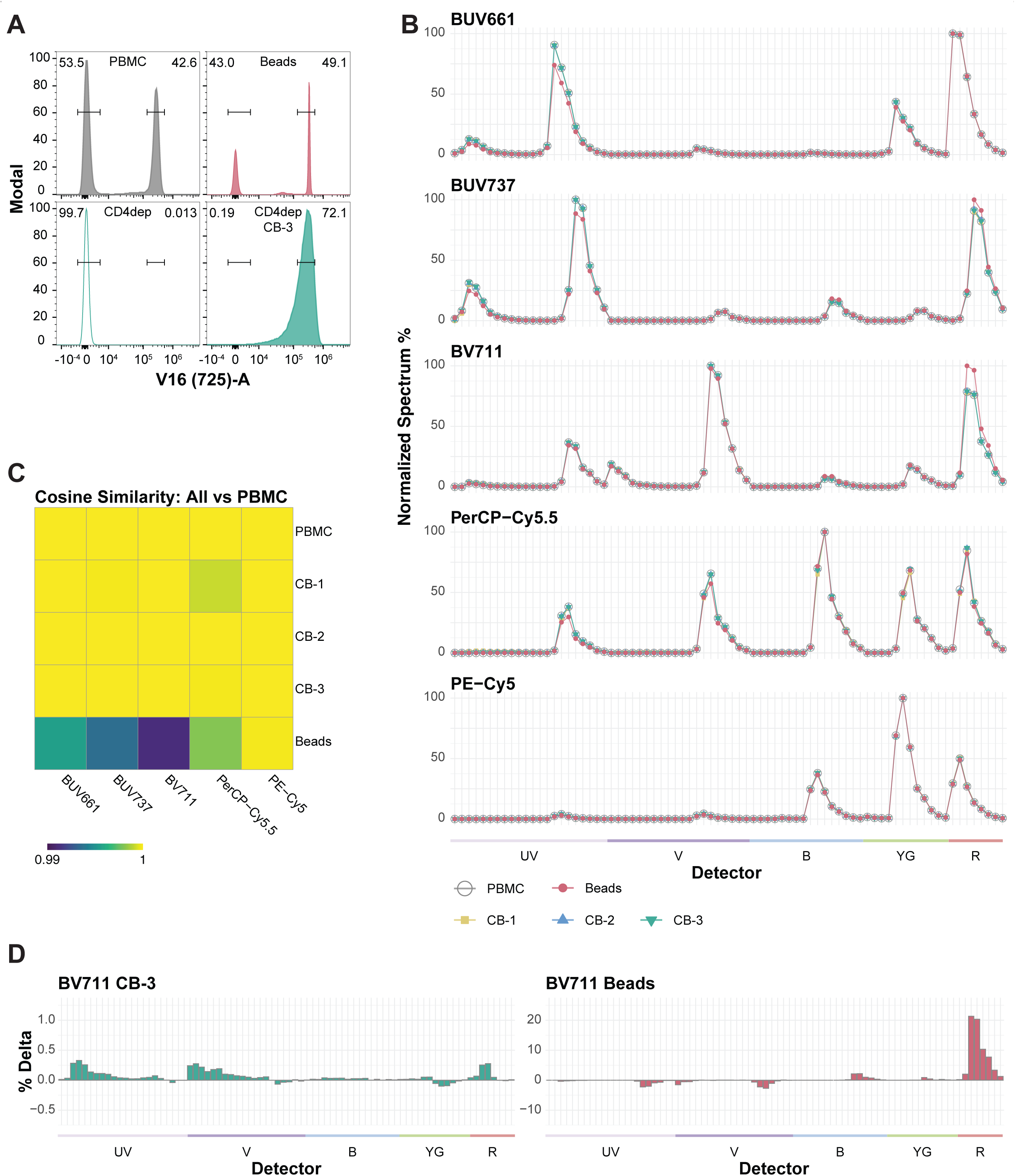
CaptureBodies recapitulate normalized fluorescence of directly stained cells. (**A**) Example of standardized flow cytometry MFI gating of lymphocytes across all groups stained with CD4 BV711 antibodies: (top left) - PBMC, (top right) - beads, (bottom left) - CD4 depleted PBMCs, and (bottom right) - CD4 depleted PBMCs with CB-3. (**B**) Overlaid normalized emission spectra for PBMC (open circle), CD4^-^ PBMC with CB-1 (yellow square), CD4^-^ PBMC with CB-2 (blue triangle), CD4^-^ PBMC with CB-3 (green triangle), or beads (red circle) across all detector channels on the BD FACSDiscover™ S8 Cell Sorter for the fluorochromes BUV661, BUV737, BV711, PerCP-Cy5.5, and PE-Cy5. (**C**) Cosine similarity for difference in spectral emission between PBMC and either each CaptureBody or compensation beads, measured across five different fluorochromes. (**D**) Visualized percent change in normalized emission spectra across all detection channels. Shown are CD4^-^ PBMC with CB-3 and beads spectra when compared to PBMC following CD4 BV711 staining. See Supplemental Figures 2-3 for all other CaptureBody formats

## Discussion

We initially addressed whether each CaptureBody could bind to mouse IgG or human CD45 separately. During BLI, CB-1, which uses a scFv format for the human CD45 binding domain, exhibited both a longer time to antigen saturation and greater dissociation for human CD45 than either CB-2 or CB-3, which use a nanobody format. Interestingly, CB-1 also exhibited both a longer time to antigen saturation and greater dissociation for mouse IgG as compared to either CB-2 or CB-3. CB-1 and CB-2 both use the nanobody clone TP1107 for the mouse Ig binding domain. We postulate the difference in IgG binding kinetics is an effect of the bispecific configuration where clone TP1107 performs optimally on the N-terminus, as is CB-2’s configuration. By BLI, CB-2 and CB-3 proved to be our favored candidates due to their low dissociation rates for either antigen which would be favorable for flow cytometry where samples are routinely acquired days after staining.

In our initial characterization, we next optimized the oligomerization of our CaptureBodies. The implementation of GCN4, a yeast transcriptional regulator, as a domain linker had a significant impact on oligomerization. GCN4 naturally forms coiled-coils because of its defining heptad repeat (*abcdefg)_n_* with hydrophobic residues at positions *a* and *d*.^34^ Harbury et al. first introduced residue modifications to drive dimer, trimer, and higher order complexes and found the Asn^16^ residue significantly drove dimer formation.^35^ Mahrenholz et al. built upon these findings and computationally predicted that the GCN4 amino acid modifications V23K and K27E would favor dimer formation.^29^ Our data demonstrate that these modifications indeed preferentially drive dimer formation. We foresee this linker optimization having broad application for designing multimer-specific antibodies as it potentially minimizes the need for downstream purification of undesirable oligomerization states while raising production yields. In respect to oligomerization, CB-2 and CB-3 were our favored candidates where after GCN4 optimization they multimerized into dimers.

Modern approaches to flow cytometry often use a combination of fluorochromes where a given fluorochrome has emission into a detector that was dedicated for a different fluorochrome. Under these circumstances, fluorescent abundances are determined by an ordinary least squares (OLS) model in the process of spectral unmixing.^36^ For accurate modeling of fluorescent abundances, single stain controls are created for each fluorescent probe to determine the relative fluorescence a probe has into each detector. These are the values used for the explanatory variables implemented in the OLS model and are referred to as the normalized spectra or spectral emission of a fluorochrome. When the normalized spectra from the single stain controls do not fully recapitulate the fluorescence observed on the full stain experimental sample for which spectral unmixing is applied to, inaccuracies can appear in the unmixed data as either false positive signal, hyper negative signal, or increased variance and ultimately lower resolution to resolve true positive signal of modeled fluorescent abundances.^5,10,11^

Traditionally, when single stain controls for conventional flow cytometry are not accurate, it is standard to address modeling errors by visual inspection of the compensated data and adjustment of the spillover coefficients (synonymous with normalized spectra). This process is labor intensive, requires expert knowledge of protein expression patterns and expected emission of fluorochromes, and works best with small panels (i.e. for a 12 fluorochrome by 12 detector spillover coefficient matrix there are 144 relationships to inspect). This is no longer advisable for data collected on full spectrum cytometers or using high dimensional panels, as the pairwise relationships between modeled fluorescent parameters and the number of detectors is much greater (i.e. for a 50 fluorochrome by 78 detector normalized spectral matrix there are 3,900 relationships to inspect). More often, a practitioner trying to manually correct modeling error in unmixed data will likely introduce random error while in pursuit of adjusting another error. Therefore, it is becoming more important for data collected on spectral cytometers or using high-parameter panels to implement accurate single stain controls to prevent modeling error from occurring in the first place.

Three guidelines have been established to generate high quality single stain controls.^37,7^ In short, these are (#1) the positive population of a single stain control must be as bright or brighter than the sample to which Compensation or Unmixing will be applied, (#2) the background fluorescence should be the same for the positive population and negative population of a single stain control, and (#3) the same fluorescent reagent used for the experimental sample should be used for the generation of the single stain control (i.e. use the exact same vial/lot of a fluorochrome-conjugated antibody). With respect to guideline (#1), we chose CD45 as the cell binding domain because CD45 has one of the highest protein densities on immune cells. It has been previously shown that PBMCs express approximately 100,000 CD45 molecules per cell (stained with anti-CD45 PE clone HI30 at saturating conditions).^38^ At saturation, PBMCs stained with CB-1, CB-2, CB-3 yielded 70-80% of the MFI that we measured when PBMCs were directly stained with anti-CD45 PE (clone HI30). This equates to approximately 70,000-80,000 antibodies bound per cell which is well above antigen binding capacity values seen for all antibody clones binding CD1 to CD100 on PBMCs.^38^ Therefore, CaptureBodies will have applicability in modeling a vast range of antigen targets.

Regarding guideline (#2), problems with single stain controls can occur when the labeled positive cell population has a unique autofluorescence that differs from the negative population. Statistical frameworks have been created to solve this problem by scatter matching and implementing a robust linear modelling approach for the generation of normalized spectra values.^3^ Our CaptureBody approach will avoid this problem since CD45 is uniformly expressed by all cells within a PBMC sample. This creates a set-up in which all stained cells are positive, while the unstained control is negative for the fluorescent conjugated antibody, thereby removing the possibility of mismatching the autofluorescence signatures observed between the two.

Regarding guideline (#3), the requirement that the identical fluorescent reagent is used for both the experimental sample and the single stain has been particularly challenging for creating single stain controls with a rare protein target or a rare cell type. In such a scenario, one could reason that in a single stain control antibody A (low cell expression or rare cell) could be replaced with an antibody B (abundantly expressed) provided that both antibodies are coupled to the same fluorochrome. However, a previous study highlighted that different lots of the same fluorochrome-conjugated antibody can produce different normalized spectra, especially when tandem fluorochromes are used.^37^ Of note, most commercially available fluorochromes for flow cytometry are tandem dyes and are extensively used in panels that range from 40^10,39,40,41,42^ to 50 colors.^5^ In these instances, compensation/unmixing particles are preferentially used but fail to reproduce the emission spectrum for certain fluorochromes when bound to cells and introduce technical errors.

We specifically compared four tandem fluorochromes that have altered normalized spectra when bound to compensation/unmixing particles to our CaptureBodies. We found that each CaptureBody reproduced the normalized spectra observed by direct staining with high similarity and low individual error for any given detector. While each CaptureBody outperformed compensation/unmixing particles for all fluorochromes tested, CB-2 and CB-3 are our favored candidates among the CaptureBodies for their improved modeling of PerCP-Cy5.5 over CB-1. Overall, CB-2 and CB-3 perform similarly, and we thus suggest that CB-3 should be suitable for capturing most mouse anti-human antibodies given its specificity for IgG(κ) while CB-2, due to it cross-species reactivity, will have use for clones generated from rat or human using an IgG1 isotype.

In summary, advancements in cytometer hardware and fluorochrome chemistry have allowed for the development of high parameters panels, however, generating precise single stain controls has remained as a significant bottleneck to further growth. We propose that using a CaptureBody will provide a systematic, reproducible, and cell-based approach to generate precise single stain controls. We expect that the CaptureBody (CB-3) will not only greatly facilitate spectral unmixing but also be immensely helpful for conventional flow cytometry and even small panels when experimental samples are limited, markers of interest are rare, or compensation/unmixing particles would generate differing spectral profiles.

## Limitations of the study

There are two notable limitations to the universal implementation of our CaptureBody reagent. CD45 was chosen as the cell-binding target for its abundant, uniform, and high antigen density in all immune cell subsets.^38^ However, large nonhematopoietic-derived cells may express target antigens at a higher frequency/density than CD45 in immune cells, therefore in these cases, CaptureBodies will not generate a bright enough single stain spectra to accurately unmix or compensate resulting flow cytometry data. Additionally, while the antibody capture targets of IgG1 and IgG(κ) in CB-2 and CB-3, respectively, will work with most antibodies, there are some fluorochrome-conjugated antibodies that do not contain these epitopes.

## Methods Section

### CaptureBody derivations

The anti-CD45 scFv was derived from the heavy and light chain variable regions of the anti-human CD45 monoclonal antibody hybridoma clone BC8^24^, which has recently been developed into the therapeutic drug for acute myeloid leukemia Apamistamab by Actinium Pharmaceuticals, Inc.^23^ The anti-CD45 nanobody was derived from the VHH variable region of a commercially available llama antibody, clone 2H5, originally described by Rokkam and Lupardus in 2020.^25,26^ These anti-CD45 components both specifically target the RO region, i.e., included in all known forms, of the human CD45 protein. Anti-mouse IgK- and IgG1 Fc-specific nanobodies were derived from alpaca VHH antibody clones 1170 and 1107, respectively, originally described by Pleiner et.al., 2018.^21^ Original GCN4 linker is a basic leucine zipper (bZIP) domain of bZIP transcription factors [a DNA-binding and dimerization domain; “cl21462,” db_xref=“CDD:473870.”] and has been used previously to create a wide variety of synthetic protein dimers constructs. Mutations for creating an optimized dimer forming GCN4 region were selected from modeling and predictions published by Mahrenholz, et.al., 2011.^29^

### Capturebody vector construction

Full length CaptureBody sequences were de novo synthesized as human codon optimized gBlock HiFi Gene Fragments by Integrated DNA Technologies (IDT, Coralville, Iowa) and cloned into the pTT3 expression vector, described by Durocher et.al.^43^, containing a Kozak consensus sequence and in-frame N-terminal murine HC signal peptide using the In-Fusion HD Cloning Kit (Takara Bio USA, Inc., Cat: 102518) per manufacturer’s protocol. pTT3 vectors with cloned CaptureBody sequences were transformed into NEB 5-alpha E. coli high efficiency competent cells (New England BioLabs, Cat: C2987H) per manufacturer’s protocol. Transformed E. coli was grown under selection for Ampicillin resistance on agar plates containing 100 ug/ml Ampicillin. Positive colonies were grown in LB broth containing 100 ug/ml Ampicillin overnight, and DNA was extracted using QIAprep Spin Miniprep or Plasmid Plus Midi Kits (Qiagen). All Capturebody clones used for expression were sequence verified by Sanger sequencing (Azenta).

### CB-1 & CB-2 Linker Optimization and CB-3 Generation

Modified CB-1 and CB-2 “Post-GCN4” Capturebody variants were constructed by substituting the original internal 6xHis-GCN4-6xHis linker with the optimized dimer-forming GCN4 sequence using QuikChange II XL Site-Directed Mutagenesis Kit per manufacturer’s protocol (Agilent, Santa Clara, CA). CB-3, containing an N-terminal anti-mouse IgK-specific 1170 nanobody and a C-terminal anti-CD45 nanobody, clone 2H5, was only constructed using the optimized post-GCN4 dimer region.

### Capturebody dimer expression and purification

Capturebodies were expressed by transient transfection of HEK293E suspension cells grown in Freestyle 293 media (ThermoFisher, Cat: 12338018) and purified by Ni-chelation chromatography followed size-exclusion chromatography (SEC) as previously described by McGuire, et.al., 2016.^44^ Briefly, HEK293E cells at a density of 10^6^ cells/mL were transfected with pTT3-capturebody constructs using 500 mg of plasmid DNA and 2 mg filter sterilized PEImax (MW 40K, pH 7.0, Polysciences, Cat: 24765) per liter of culture. Expression was carried out for 6 days at 37°C and 5% CO2 in shaker flasks (@ 125 rpm), after which cells and cellular debris were removed by centrifugation at 4,000 × g followed by filtration through a 0.22 μm filter. Clarified supernatant was adjusted to 2 mM Imidazole and incubated with His60 Ni-Superflow resin (Takara Bio, Cat: 635660) overnight at 4°C followed by extensive washing with 0.15 M NaCl, 20 mM Tris, 5 mM imidazole (pH 8.0), and then eluted with 250 mM imidazole, 0.3 M NaCl, 20 mM Tris (pH 8.0). Affinity purified protein was concentrated using an Amicon Ultra-15 centrifugal filter with 10kDa MWCO (MilliporeSigma, Cat: UFC9010), clarified over a 0.22 mm filter (Millipore, Cat: UFC30GV00), and then purified on an AKTA pure system (General Electric) using a Superdex 200 increase 10/300 column (Cytiva, Cat: 28990944) into Tris buffer (pH 7.5). SEC fractions containing dimer or trimer peaks were collected, pooled separately, and stored at 4°C.

### Biolayer interferometry and protein gel electrophoresis

Capturebodies were evaluated for bi-specific antigen recognition by biolayer interferometry essentially as described previously by Gray, et.al., 2022,^45^ using the Octet Red96 instrument (Sartorius, Göttingen, Germany) at 29°C with shaking at 500 rpm. To estimate binding of Capturebody dimer anti-mouse IgG Fc and anti-mouse IgG(κ) specific domains, total purified mouse IgG was immobilized on anti-mouse Fc (AMC, Sartorius, Cat: 18-5088) biosensors (@ 40 μg/ml in PBS) for 240s. To estimate binding of Capturebody dimer anti-human CD45 specific domains, a recombinant human CD45 protein containing a human Fc tag (Sino Biological, Inc., Cat: 10086-H02H) was loaded onto anti-human Fc (AHC, Sartorius, Cat: 18-5063) biosensors (@ 1 uM diluted in PBS) for 240s. Antigen loaded biosensors were then incubated for 1 min in PBS to establish baseline signals. Mouse IgG and human CD45 loaded sensors were then dipped into clarified conditioned media from freshly harvested Capturebody expression cultures for 300 seconds to observe the association, followed by immersion in PBS alone for 300 seconds to measure potential dissociation. All measurements of Capturebody binding were corrected by subtracting the background signal obtained from duplicate traces generated with the same loaded antigen but dipped into PBS only. SEC purified Capturebody dimers (10 ug per well) were subjected to SDS-gel electrophoresis using NuPAGE Bis-Tris Mini Protein Gels (4–12%, ThermoFisher, Cat: NP0321BOX) in 1x MES buffer and 3 ul PageRule Plus Prestained Protein Ladder was used as molecular weight standard (10 to 250 kDa, ThermoFisher, Cat: 26619). Separated proteins were fixed and visualized using 1% Coomassie Brilliant blue stain G 250 (Bio-Rad, Cat: 20279) in 40% Methanol plus 10% Acetic Acid and destained gels were photographed on a ChemiDoc Go Imaging System (Bio-Rad).

### Primary Human Samples and Processing

Whole blood or leukopak were purchased from Bloodworks Northwest (https://www.bloodworksnw.org/, Washington, USA).

Whole blood (200 mL) was diluted 1:1 with PBS (Gibco, Cat: 14190-144) containing 2% FBS (Peak Serum, Cat: PS-FBa2) and then processed using SepMate tubes (StemCell Technologies, Cat: 85450) and Lymphoprep (Stem Cell Technologies, Cat: 07851) according to manufacturer protocols. In brief, diluted blood layered on Lymphoprep in SepMate tubes were centrifuged for 1200 x g for 16 minutes. After centrifugation, the supernatant above the mononuclear cell layer was pipetted and discarded to reduce platelet contamination. The remaining contents were then poured off into a new tube. Samples were then centrifuged for 400 x g for 5 minutes, decanted, and resuspended in PBS containing 2% FBS. As a second measure to reduce platelets, samples were centrifuged for 120 x g for 10 minutes, decanted, and resuspended in PBS containing 2% FBS.

Leukopak were processed using the protocol outlined by StemCell Technologies (https://www.stemcell.com/leukopak-processing-protocol.html). To wash, leukopak contents were aliquoted, diluted 1:1 with PBS containing 2% FBS, centrifuged for 300 x g for 10 minutes, and the supernatant decanted. To lyse red blood cells, the pellet was resuspended in ACK Lysing Buffer (Gibco, Cat: A10492-01) at a 4:1 (lysis solution to sample) volume ratio and incubated at room temperature for 5 minutes. Samples were then topped up to 50 mL with PBS containing 2% FBS, centrifuged for 300 x g for 10 minutes, and the supernatant decanted. Red blood cell lysis was repeated for a total of two lysis steps. To reduce platelets, samples were resuspended in PBS containing 2% FBS were centrifuged for 120 x g for 10 minutes, decanted, and resuspended in PBS containing 2% FBS. Platelet removal was repeated once for a total of two platelet removal steps.

Samples from either process were counted using Trypan Blue Stain (0.4%) (Gibco, Cat: 15250-061) and a TC20 Automated Cell Counter (Bio-Rad Laboratories). For long term storage, samples were resuspended in Cell Recovery Freezing Media (Gibco, Cat: 12648-010) or 90 % FBS with 10% DMSO (Sigma, Cat: D2650-5X 10ML), placed in a Stratacooler (Agilent, Cat: 400005) at -80 °C for 1-3 days, and then transferred to liquid nitrogen storage.

### Titration of CaptureBody reagent

After thawing, PBMCs were incubated for 30 minutes with each CaptureBody starting at a max concentration of 0.5 µg and continuing with 11 two-fold serial dilutions down to 0.244 ng in a 50 µL volume of PBS. Cells were then washed twice; 150-200 µL of PBS was added to cells, centrifuged for 400 x g for 5 minutes, and then the supernatant decanted. Cells were then incubated for 30 minutes with 0.25 µg of PE Isotype Control Antibody (Clone: MOPC-21, Mouse IgG1 κ) in a 50 µL volume of PBS. Cells were then washed twice; 150-200 µL of PBS was added to cells, centrifuged for 400 x g for 5 minutes, and then the supernatant decanted. To fix cells, 100 µL of Fixation/Permeabilization solution (BD Biosciences, Cat: 51-2090KZ) was added to cells and incubated for 15 minutes. Cells were then washed twice; 100-200 µL of PBS was added to cells, centrifuged for 800 x g for 5 minutes, and then the supernatant decanted. Cells were resuspended in 200 µL of PBS and stored at 4 °C in the dark until acquisition.

### Comparison of CaptureBody staining to direct anti-CD45 staining of cells

After thawing, PBMCs were incubated for 30 minutes with each CaptureBody at a concentration of 0.5 µg in a 50 µL volume of PBS or in PBS alone. Cells were then washed twice; 150-200 µL of PBS was added to cells, centrifuged for 400 x g for 5 minutes, and then the supernatant decanted. To the cells incubated with a Capture Body, cells were then incubated with 0.25 µg of PE Isotype Control Antibody (Clone: MOPC-21, Mouse IgG1 κ) in a 50 µL volume of PBS. To the cells prior incubated in PBS alone, cells were then incubated with 0.25 µg of PE anti-CD45 (Clone: HI30 Mouse IgG1 κ) in a 50 µL volume of PBS, 0.25 µg of PE Mouse IgG1 κ Isotype Control Antibody (Clone: MOPC-21, Mouse IgG1 κ) in a 50 µL volume of PBS, or PBS alone. Cells were then washed twice; 150-200 µL of PBS was added to cells, centrifuged for 400 x g for 5 minutes, and then the supernatant decanted. To fix cells, 100 µL of Fixation/Permeabilization solution was added to cells and incubated for 15 minutes. Cells were then washed twice; 100-200 µL of PBS was added to cells, centrifuged for 800 x g for 5 minutes, and then the supernatant decanted. Cells were resuspended in 200 µL of PBS and stored at 4 °C in the dark until acquisition.

### Generation of single stain controls

After thawing PBMCs, CD4 depleted PBMCs were generated using the Human Biotin Selection Kit (StemCell Technologies, Cat: 18553) per the manufacturer’s recommended protocol, but without blocking Fc receptors (step adding in anti-CD32) to not add in Ig that may interact with CaptureBodies. For the depletion protocol, Biotin anti-CD4 (Clone: SK3, Mouse IgG1 κ) was used at 5 µg/mL concentration per sample.

CD4 depleted PBMCs were incubated for 30 minutes with each CaptureBody at a concentration of 0.5 µg in a 50 µL volume of PBS or in PBS alone. Additionally at this step, unfractionated PBMCs were only incubated in 50 µL of PBS alone. Cells were then washed twice; 150-200 µL of PBS was added to cells, centrifuged for 400 x g for 5 minutes, and then the supernatant decanted. Then for each condition, 0.125 µg of anti-CD4 (Clone: SK3, Mouse IgG1 κ) conjugated to BUV661, BUV737, BV711, PerCP-Cy5.5, or PE-Cy5 was added in a 50 µL volume of PBS and incubated for 30 minutes. Cells were then washed twice; 150-200 µL of PBS was added to cells, centrifuged for 400 x g for 5 minutes, and then the supernatant decanted. To fix cells, 100 µL of Fixation/Permeabilization solution was added to cells and incubated for 15 minutes. Cells were then washed twice; 100-200 µL of PBS was added to cells, centrifuged for 800 x g for 5 minutes, and then the supernatant decanted. Cells were resuspended in 200 µL of PBS and stored at 4 °C in the dark until acquisition. Single stain controls were additionally made with a polystyrene based compensation particle, CompBeads Anti-Mouse Ig κ (BD, Cat: 552843), prepared using the exact same anti-CD4 conjugated fluorochromes and treated experimentally the same as cell based single stain controls.

### Acquisition of flow cytometry samples

All samples were acquired on a single FACSDiscover S8 Cell Sorter (BD Biosciences). The cytometer contains 78 fluorescent APD detectors and 5 lasers: 349 nm (30 mW), 405 nm (50 mW), 488 nm (100 mW), 561 nm (50 mW), and 637 nm (100 mW). The cytometer uses FACSChorus software.

For experiments determining the normalized emission spectrum of single stain controls (data in Figures 4), detector gains were used based on the manufacturer’s daily set up and quality control. For titration of CaptureBodies using PE Isotype Control Antibody (Clone: MOPC-21, Mouse IgG1 κ) and comparison of CaptureBodies to PE anti-CD45 (Clone: HI30 Mouse IgG1 κ) (data in Figures 3), detector gains were lowered from the manufacturer’s daily set up and quality control settings for only the yellow-green laser detector array so that PE anti-CD45 stained PBMCs were just on scale for the primary detector YG1 (575-A). The lowered detectors settings were held constant for all experiments using PE.

Experiment files were exported as FCS3.2 format from FACSChorus. FlowJo (version 11.1.0) was used to determine the normalized emission spectrum for each fluorochrome with Autospill turned off. Positive gate thresholds were kept constant for all samples of a particular fluorochrome so that the MFI of the positive population would be equivalent between different samples when generating the normalized emission spectrum. FlowJo (version 10.10.0) was used gate, generate flow plots, and determine all other tabular statistics for flow cytometry data.

### Statistics and graphical representation

R was used to graph and process tabular flow cytometry data. The following libraries were used to read and organize data; tidyr, dyplyr, readr, and stringr. Cosine similarity was computed using the cosine function from the lsa package. pheatmap, viridis, RcolorBrewer, and ggplot2 were used for plotting. Graphs were formatted for publication using Adobe Illustrator.

### AlphaFold3 Structure Prediction and Region Delineation

CaptureBody amino acid sequences were formatted into JSON files and submitted to AlphaFold3 locally on Fred Hutch servers. One seed was run per CaptureBody, generating five diffusion samples, each ranked from lowest to highest ranking_score. The diffusion sample with the highest ranking_score was then used as the final structure output. The resulting output was visualized in UCSF ChimeraX^46^ and specific regions were delineated using hexadecimal codes: CD45 scFv heavy chain - #FA8072, CD45 scFv light chain #FFDAB9, scFv linker - #808080, CD45 nanobody - #FFD700, IgG1-Fc nanobody - #FF8C00, IgG(k) nanobody - #00BFFF, GCN4 linker and HisTag - #F0F8FF.

## Acknowledgments

This research was supported by the Flow Cytometry Shared Resource (RRID:SCR_022613) of the Fred Hutch/University of Washington/Seattle Children’s Cancer Consortium (P30 CA015704). We would like to thank Michele Black for her state-of-the-art flow cytometry expertise and exemplary leadership in running the Flow Cytometry Shared Resource. This work is the result of NIH funding, in whole or in part, and is subject to the NIH Public Access Policy. Through acceptance of this federal funding, the NIH has been given a right to make the work publicly available in PubMed Central. This work was supported by NIH grants R01AI123323, R01AI179712 and 1R56DE032009 (to M.P.). The graphical abstract and Figure 1 were created with BioRender.

## Author Contributions

Conceptualization: A.K. and M.P, Methodology: A.K., M.G., Investigation: A.Z., A.K., M.G., L.K.S., Formal analysis: A.Z., A.K., M.G., M.P. Visualization: A.Z., A.K., M.G., M.P. Writing, original draft: A.Z., Writing, review, and editing: A.Z., A.K., M.G., M.P., Funding: M.P., Supervision: M.P.

## Declaration of Interests

The authors declare no competing interests.

## Notes

### Competing Interest Statement

The authors have declared no competing interest.

## References

1. Nolan, J.P. (2022). The evolution of spectral flow cytometry. Cytometry A 101, 812–817. 10.1002/cyto.a.24566.

2. Astakhova, E.A., Gubaeva, A.S., Naumova, D.A., Egorova, A.E., Maznina, A.A., Rybkina, I.G., Osmanov, I.M., Tabakov, D.V., Mityaeva, O.N., and Volchkov, P.Y. (2025). Spectral Flow Cytometry: The Current State and Future of the Technology. Int J Mol Sci 26. 10.3390/ijms26125911.

3. Burton, O.T., Bücken, L., De Vuyst, L., Humblet-Baron, S., De Leon, A.L.M., Khan, S., Cerveira, J., Dooley, J., and Liston, A. (2025). AutoSpectral improves spectral flow cytometry accuracy through optimised spectral unmixing and autofluorescence-matching at the cellular level. bioRxiv, 2025.2010.2027.684855. 10.1101/2025.10.27.684855.

4. Kmet, R., and Novo, D. (2025). Reducing Spreading: Removing the Impact of Irrelevant Dyes Improves Unmixed Flow Cytometry Data. Cytometry A 107, 573–586. 10.1002/cyto.a.24957.

5. Konecny, A.J., Mage, P.L., Tyznik, A.J., Prlic, M., and Mair, F. (2024). OMIP-102: 50-color phenotyping of the human immune system with in-depth assessment of T cells and dendritic cells. Cytometry A 105, 430–436. 10.1002/cyto.a.24841.

6. Ferrer-Font, L., Pellefigues, C., Mayer, J.U., Small, S.J., Jaimes, M.C., and Price, K.M. (2020). Panel Design and Optimization for High-Dimensional Immunophenotyping Assays Using Spectral Flow Cytometry. Curr Protoc Cytom 92, e70. 10.1002/cpcy.70.

7. Ferrer-Font, L., Small, S.J., Lewer, B., Pilkington, K.R., Johnston, L.K., Park, L.M., Lannigan, J., Jaimes, M.C., and Price, K.M. (2021). Panel Optimization for High-Dimensional Immunophenotyping Assays Using Full-Spectrum Flow Cytometry. Curr Protoc 1, e222. 10.1002/cpz1.222.

8. Futamura, K., Sekino, M., Hata, A., Ikebuchi, R., Nakanishi, Y., Egawa, G., Kabashima, K., Watanabe, T., Furuki, M., and Tomura, M. (2015). Novel full-spectral flow cytometry with multiple spectrally-adjacent fluorescent proteins and fluorochromes and visualization of in vivo cellular movement. Cytometry A 87, 830–842. 10.1002/cyto.a.22725.

9. Novo, D. (2022). A comparison of spectral unmixing to conventional compensation for the calculation of fluorochrome abundances from flow cytometric data. Cytometry A 101, 885–891. 10.1002/cyto.a.24669.

10. Park, L.M., Lannigan, J., and Jaimes, M.C. (2020). OMIP-069: Forty-Color Full Spectrum Flow Cytometry Panel for Deep Immunophenotyping of Major Cell Subsets in Human Peripheral Blood. Cytometry A 97, 1044–1051. 10.1002/cyto.a.24213.

11. Shevchenko, Y., Lurje, I., Tacke, F., and Hammerich, L. (2024). Fluorochrome - dependent specific changes in spectral profiles using different compensation beads or primary cells in full spectrum cytometry. Cytometry Part A 105, 458–463. 10.1002/cyto.a.24836.

12. Brinkmann, U., and Kontermann, R.E. (2017). The making of bispecific antibodies. MAbs 9, 182–212. 10.1080/19420862.2016.1268307.

13. Labrijn, A.F., Janmaat, M.L., Reichert, J.M., and Parren, P. (2019). Bispecific antibodies: a mechanistic review of the pipeline. Nat Rev Drug Discov 18, 585–608. 10.1038/s41573-019-0028-1.

14. Sun, Y., Yu, X., Wang, X., Yuan, K., Wang, G., Hu, L., Zhang, G., Pei, W., Wang, L., Sun, C., and Yang, P. (2023). Bispecific antibodies in cancer therapy: Target selection and regulatory requirements. Acta Pharm Sin B 13, 3583–3597. 10.1016/j.apsb.2023.05.023.

15. Reiter, Y., Brinkmann, U., Kreitman, R.J., Jung, S.H., Lee, B., and Pastan, I. (1994). Stabilization of the Fv fragments in recombinant immunotoxins by disulfide bonds engineered into conserved framework regions. Biochemistry 33, 5451–5459. 10.1021/bi00184a014.

16. Vincke, C., and Muyldermans, S. (2012). Introduction to heavy chain antibodies and derived Nanobodies. Methods Mol Biol 911, 15–26. 10.1007/978-1-61779-968-6_2.

17. Alexander, E., and Leong, K.W. (2024). Discovery of nanobodies: a comprehensive review of their applications and potential over the past five years. J Nanobiotechnology 22, 661. 10.1186/s12951-024-02900-y.

18. Muyldermans, S. (2013). Nanobodies: natural single-domain antibodies. Annu Rev Biochem 82, 775–797. 10.1146/annurev-biochem-063011-092449.

19. Hambach, J., Mann, A.M., Bannas, P., and Koch-Nolte, F. (2022). Targeting multiple myeloma with nanobody-based heavy chain antibodies, bispecific killer cell engagers, chimeric antigen receptors, and nanobody-displaying AAV vectors. Front Immunol 13, 1005800. 10.3389/fimmu.2022.1005800.

20. Arbabi Ghahroudi, M., Desmyter, A., Wyns, L., Hamers, R., and Muyldermans, S. (1997). Selection and identification of single domain antibody fragments from camel heavy-chain antibodies. FEBS Lett 414, 521–526. 10.1016/s0014-5793(97)01062-4.

21. Pleiner, T., Bates, M., and Gorlich, D. (2018). A toolbox of anti-mouse and anti-rabbit IgG secondary nanobodies. J Cell Biol 217, 1143–1154. 10.1083/jcb.201709115.

22. Boitano, A., Cooke, M., Gillard, G.O., McDonagh, C.F., Palchaudhuri, R., Pearse, B.R. (2022) ANTI-CD45 ANTIBODY DRUG CONJUGATES AND USES THEREOF. patent U.S. Patent Application Publication No. US 2022/0175951 A1.

23. Gyurkocza, B., Nath, R., Seropian, S., Choe, H., Litzow, M.R., Abboud, C., Koshy, N., Stiff, P., Tomlinson, B., Abhyankar, S., et al. (2025). Randomized Phase III SIERRA Trial of (131)I-Apamistamab Before Allogeneic Hematopoietic Cell Transplantation Versus Conventional Care for Relapsed/Refractory AML. J Clin Oncol 43, 201–213. 10.1200/JCO.23.02018.

24. Matthews, D.C., Appelbaum, F.R., Eary, J.F., Fisher, D.R., Durack, L.D., Hui, T.E., Martin, P.J., Mitchell, D., Press, O.W., Storb, R., and Bernstein, I.D. (1999). Phase I study of (131)I-anti-CD45 antibody plus cyclophosphamide and total body irradiation for advanced acute leukemia and myelodysplastic syndrome. Blood 94, 1237–1247.

25. Lupardus, P.J., Rokkam, D. (2023) CD45 BINDING MOLECULES AND METHODS OF USE. patent U.S. Patent Application Publication No. US 2023/0357424 A1.

26. Rokkam, D., and Lupardus, P.J. (2020). Discovery and characterization of llama VHH targeting the RO form of human CD45. bioRxiv, 2020.2009.2001.278853. 10.1101/2020.09.01.278853.

27. McKenney, D.W., Onodera, H., Gorman, L., Mimura, T., and Rothstein, D.M. (1995). Distinct isoforms of the CD45 protein-tyrosine phosphatase differentially regulate interleukin 2 secretion and activation signal pathways involving Vav in T cells. J Biol Chem 270, 24949–24954. 10.1074/jbc.270.42.24949.

28. McNeill, L., Cassady, R.L., Sarkardei, S., Cooper, J.C., Morgan, G., and Alexander, D.R. (2004). CD45 isoforms in T cell signalling and development. Immunol Lett 92, 125–134. 10.1016/j.imlet.2003.10.018.

29. Mahrenholz, C.C., Abfalter, I.G., Bodenhofer, U., Volkmer, R., and Hochreiter, S. (2011). Complex networks govern coiled-coil oligomerization--predicting and profiling by means of a machine learning approach. Mol Cell Proteomics 10, M110 004994. 10.1074/mcp.M110.004994.

30. Holmes, K.L., and Lantz, L.M. (2001). Chapter 9 Protein labeling with fluorescent probes. In Methods in Cell Biology, (Academic Press), pp. 185–204. 10.1016/S0091-679X(01)63013-9.

31. Pannu, K.K., Joe, E.T., and Iyer, S.B. (2001). Performance evaluation of QuantiBRITE phycoerythrin beads. Cytometry 45, 250–258. 10.1002/1097-0320(20011201)45:4<250::aid-cyto10021>3.0.co;2-t.

32. Rajwa, B., and Roederer, M. (2025). Lost in Translation: Harmonizing Terminology and Defining Mathematical Tools for Panel Optimization. Cytometry A 107, 793–816. 10.1002/cyto.a.24972.

33. Kotu, V., and Deshpande, B. (2018). Data Science: Concepts and Practice (Morgan Kaufmann).

34. O’Shea, E.K., Klemm, J.D., Kim, P.S., and Alber, T. (1991). X-ray structure of the GCN4 leucine zipper, a two-stranded, parallel coiled coil. Science 254, 539–544. 10.1126/science.1948029.

35. Harbury, P.B., Zhang, T., Kim, P.S., and Alber, T. (1993). A switch between two-, three-, and four-stranded coiled coils in GCN4 leucine zipper mutants. Science 262, 1401–1407. 10.1126/science.8248779.

36. Nolan, J.P., and Condello, D. (2013). Spectral flow cytometry. Curr Protoc Cytom *Chapter 1*, 1 27 21-21 27 13. 10.1002/0471142956.cy0127s63.

37. Roederer, M. (2002). Compensation in flow cytometry. Curr Protoc Cytom *Chapter 1*, Unit 1 14. 10.1002/0471142956.cy0114s22.

38. Kalina, T., Fiser, K., Perez-Andres, M., Kuzilkova, D., Cuenca, M., Bartol, S.J.W., Blanco, E., Engel, P., and van Zelm, M.C. (2019). CD Maps-Dynamic Profiling of CD1-CD100 Surface Expression on Human Leukocyte and Lymphocyte Subsets. Front Immunol 10, 2434. 10.3389/fimmu.2019.02434.

39. Brandi, J., Wiethe, C., Riehn, M., and Jacobs, T. (2023). OMIP-93: A 41-color high parameter panel to characterize various co-inhibitory molecules and their ligands in the lymphoid and myeloid compartment in mice. Cytometry A 103, 624–630. 10.1002/cyto.a.24740.

40. Kare, A.J., Nichols, L., Zermeno, R., Raie, M.N., Tumbale, S.K., and Ferrara, K.W. (2023). OMIP-095: 40-Color spectral flow cytometry delineates all major leukocyte populations in murine lymphoid tissues. Cytometry A 103, 839–850. 10.1002/cyto.a.24788.

41. Park, L.M., Lannigan, J., Low, Q., Jaimes, M.C., and Bonilla, D.L. (2024). OMIP-109: 45-color full spectrum flow cytometry panel for deep immunophenotyping of the major lineages present in human peripheral blood mononuclear cells with emphasis on the T cell memory compartment. Cytometry A 105, 807–815. 10.1002/cyto.a.24900.

42. Waaijer, L.A., van Cranenbroek, B., and Koenen, H. (2025). OMIP-112: 42-Parameter (40-Color) Spectral Flow Cytometry Panel for Comprehensive Immunophenotyping of Human Peripheral Blood Leukocytes. Cytometry A 107, 226–232. 10.1002/cyto.a.24927.

43. Durocher, Y., Perret, S., and Kamen, A. (2002). High-level and high-throughput recombinant protein production by transient transfection of suspension-growing human 293-EBNA1 cells. Nucleic Acids Res 30, E9. 10.1093/nar/30.2.e9.

44. McGuire, A.T., Gray, M.D., Dosenovic, P., Gitlin, A.D., Freund, N.T., Petersen, J., Correnti, C., Johnsen, W., Kegel, R., Stuart, A.B., et al. (2016). Specifically modified Env immunogens activate B-cell precursors of broadly neutralizing HIV-1 antibodies in transgenic mice. Nat Commun 7, 10618. 10.1038/ncomms10618.

45. Gray, M.D., Feng, J., Weidle, C.E., Cohen, K.W., Ballweber-Fleming, L., MacCamy, A.J., Huynh, C.N., Trichka, J.J., Montefiori, D., Ferrari, G., et al. (2022). Characterization of a vaccine-elicited human antibody with sequence homology to VRC01-class antibodies that binds the C1C2 gp120 domain. Sci Adv 8, eabm3948. 10.1126/sciadv.abm3948.

46. Meng, E.C., Goddard, T.D., Pettersen, E.F., Couch, G.S., Pearson, Z.J., Morris, J.H., and Ferrin, T.E. (2023). UCSF ChimeraX: Tools for structure building and analysis. Protein Sci 32, e4792. 10.1002/pro.4792.

