## Supplemental Figures 1-3 & Tables 1-3 for "CaptureBody – an anti-CD45 x anti-IgG bispecific antibody enables accurate unmixing for spectral flow cytometry"

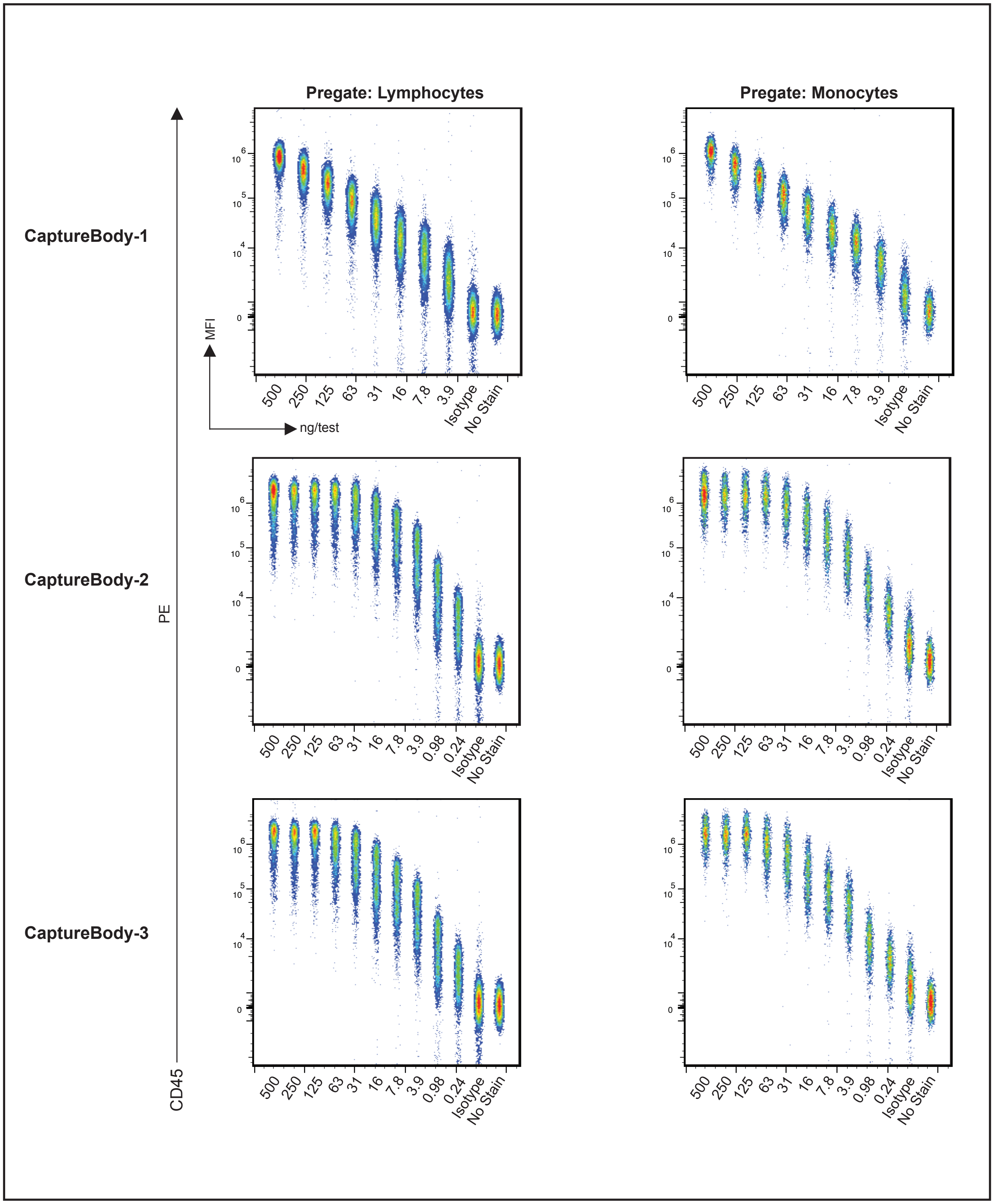

**Supplementary Figure 1: CaptureBody titer across lymphocytes and monocytes**
Antibody titrations for gated lymphocyte and monocyte MFI with each CaptureBody. Titer began at 500ng of CaptureBody per well, then underwent 2-fold dilutions from left to right. All wells were seeded with 1.0 x 10^6 PBMCs. Fluorescence was evaluated using a PE mouse IgG1 isotype antibody

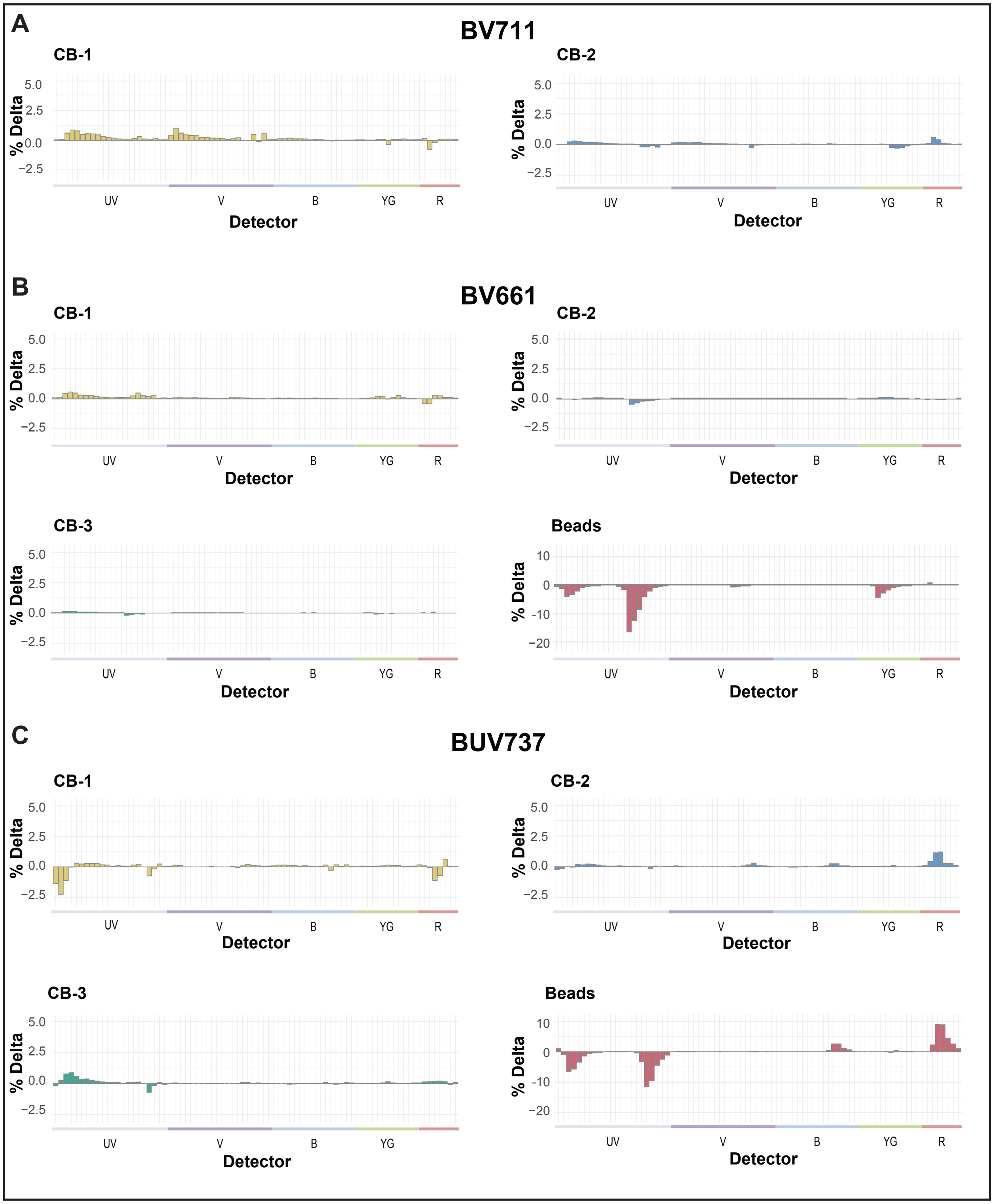

**Supplementary Figure 2: Percent change of normalized spectra per detector for Capture-Bodies or compensation/unmixing particles compared to directly stained PBMCs**

(A) Percent change of normalized spectral emission to CD4 stained PBMCs across CD4 depleted PBMCs with CB-1 & CB-2 with the CD4 fluorochrome BV711. (B) Percent change of normalized spectral emission to CD4 stained PBMCs across CD4 depleted PBMCs with CB-1, CB-2, CB-3, or only beads with the CD4 fluorochrome BV661. (C) Percent change of normalized spectral emission to CD4 stained PBMCs across CD4 depleted PBMCs with CB-1, CB-2, CB-3, or only beads with the CD4 fluorochrome BUV737

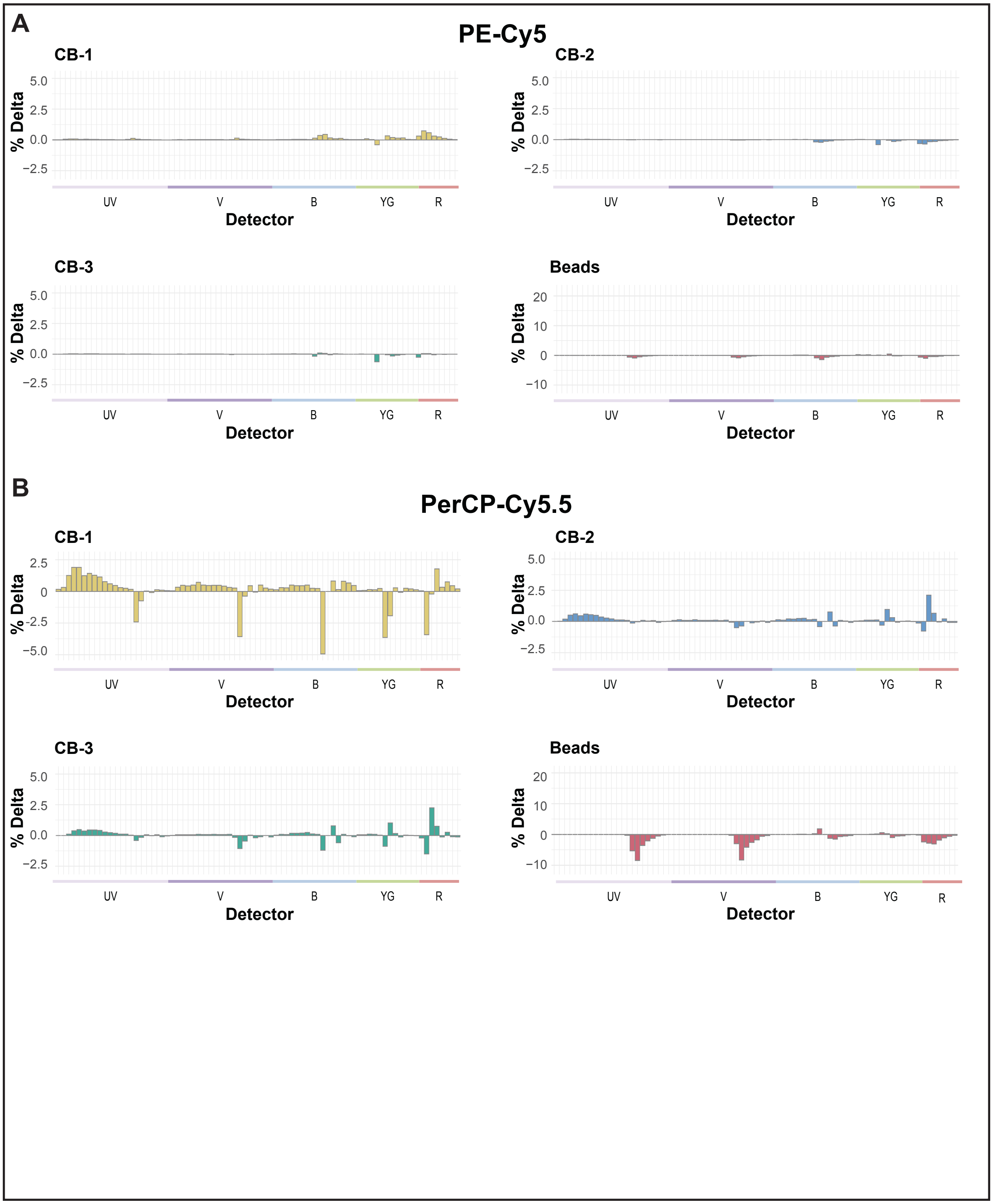

**Supplementary Figure 3: Percent change of normalized spectra per detector for CaptureBodies or compensation/unmixing particles compared to directly stained PBMCs**

(A) Percent change of normalized spectral emission to CD4 stained PBMCs across CD4 depleted PBMCs with CB-1, CB-2, CB-3, or only beads with the CD4 fluorochrome PE-Cy5
(B) Percent change of normalized spectral emission to CD4 stained PBMCs across CD4 depleted PBMCs with CB-1, CB-2, CB-3, or only beads with the CD4 fluorochrome PerCP-Cy5.5

| Name | Antigen Target | Human Codon Optimized DNA Fragment | Mature Protein Amino Acid Sequence | Amino Acid Position |
|---|---|---|---|---|
| **CaptureBody-1** | α-CD45 x α-IgG1 Fc | CATCCTTTTTCTAGTAGCAACTGCAACCGGTGTACATTCCGATATCGCTCTGACCCAGTCCCCCGCCAGCCTGGCCGTGAGCCTGGGACAGAGGGCTACCATCAGCTGCAGAGCATCCAAATCCGTGTCTACAAGCGGCTACAGCTACCTGCACTGGTACCAGCAGAAGCCTGGACAGCCACCAAAACTGCTGATCTACCTGGCCAGCAACCTGGAAAGCGGCGTGCCCGCCCGGTTCAGCGGCTCCGGCAGCGGAACTGACTTTACCCTGAACATTCACCCTGTGGAGGAGGAGGACGCCGCTACCTACTACTGCCAGCATAGTCGGGAGCTGCCATTCACATTTGGCTCCGGCACAAAGCTGGAGATTAAGGGTACAGGAGGTAGTGGCGGTGGCGGATCAGGGGGAGGAGGATCTGGCGGGGGTGCAAGCGAGGTGAAACTGCTCGAGTCCGGCGGCGGCCTGGTGCAGCCCGGCGGTTCCCTGAAGCTGTCTTGCGCCGCCTCCGGCTTCGACTTCAGCAGATATTGGATGTCATGGGTGAGGCAGGCACCCGGCAAAGGACTGGAATGGATCGGCGAAATCAACCCAACCTCTAGCACCATTAACTTCACTCCTAGCCTGAAGGATAAGGTGTTCATCTCCAGGGACAACGCCAAGAATACACTGTACCTGCAGATGAGCAAGGTGAGATCTGAGGACACCGCCCTGTATTACTGTGCCAGAGGCAACTACTACAGGTATGGCGACGCCATGGACTACTGGGGCCAGGGCACATCAGTGACCGTGAGCTCCCATCATCACCATCACCACGGAGGTTCCGGTATGAAGCAGCTGGAGGACAAGGTCGAGGAGCTGCTGAGCAAGAACTACCACCTGGAGAATGAGAAGGCCCGCCTGGAGAAGCTGGTAGGCGAGAGGGGCGGATCCCACCATCACCATCACCATGCTGGTGGGCAGGTGCAGCTGGTGGAGTCCGGGGGAGGACTGGTGCAGCCTGGCGGCAGCCTGCGGCTGAGCTGTGCCGCCTCAGGCTTCACCTTTTCCGACACTTGGATGAATTGGGTGAGACAGGCACCCGGCAAAGGCCTGTATTGGATCAGCGCCATCAACCCTGACGGCGGAAATACCGCCTACGCCGATTCCGTGAAGGGAAGATTCACTATTAGCAGGGACAATGCCAAGAACATGGTGTATCTGCAGATGGACAATCTGCGCCCCGAGGACACCGCTATGTACTACTGTGCCAAGGGATGGGTGAGACTGCCTGATCCAGACCTGGTGAGAGGCCAGGGCACCCAGGTGACCGTGTCAAGCTGATAGGGATCCTGTCGTCGGATCTCCCCCGACCTC | DIALTQSPASLAVSLGQRATISCRASKSVSTSGYSYLHWYQQKPGQPPKLLIYLASNLESGVPARFSGSGSGTDFTLNIHPVEEEDAATYYCQHSRELPFTFGSGTKLEIKGTGGSGGGGSGGGGSGGGASEVKLLESGGGLVQPGGSLKLSCAASGFDFSRYWMSWVRQAPGKGLEWIGEINPTSSTINFTPSLKDKVFISRDNAKNTLYLQMSKVRSEDTALYYCARGNYYRYGDAMDYWGQGTSVTVSSHHHHHHGGSGMKQLEDKVEELLSKNYHLENEKARLEKLVGERGGSHHHHHHAGGQVQLVESGGGLVQPGGSLRLSCAASGFTFSDTWMNWVRQAPGKGLYWISAINPDGGNTAYADSVKGRFTISRDNAKNMVYLQMDNLRPEDTAMYYCAKGWVRLPDPDLVRGQGTQVTVSS | D1-S426 |
| Mouse Ig heavy chain leader sequence | N/A | CATCCTTTTTCTAGTAGCAACTGCAACCGGTGTACATTCC | N/A | N/A |
| Apamistamab light chain variable region | α-CD45 | GATATCGCTCTGACCCAGTCCCCCGCCAGCCTGGCCGTGAGCCTGGGACAGAGGGCTACCATCAGCTGCAGAGCATCCAAATCCGTGTCTACAAGCGGCTACAGCTACCTGCACTGGTACCAGCAGAAGCCTGGACAGCCACCAAAACTGCTGATCTACCTGGCCAGCAACCTGGAAAGCGGCGTGCCCGCCCGGTTCAGCGGCTCCGGCAGCGGAACTGACTTTACCCTGAACATTCACCCTGTGGAGGAGGAGGACGCCGCTACCTACTACTGCCAGCATAGTCGGGAGCTGCCATTCACATTTGGCTCCGGCACAAAGCTGGAGATTAAG | DIALTQSPASLAVSLGQRATISCRASKSVSTSGYSYLHWYQQKPGQPPKLLIYLASNLESGVPARFSGSGSGTDFTLNIHPVEEEDAATYYCQHSRELPFTFGSGTKLEIK | D1-K111 |
| $G_4S$ scFv linker | N/A | GGTACAGGAGGTAGTGGCGGTGGCGGATCAGGGGGAGGAGGATCTGGCGGGGGTGCAAGC | GTGGSGGGGSGGGGSGGGAS | G112-S131 |
| Apamistamab heavy chain variable region | α-CD45 | GAGGTGAAACTGCTCGAGTCCGGCGGCGGCCTGGTGCAGCCCGGCGGTTCCCTGAAGCTGTCTTGCGCCGCCTCCGGCTTCGACTTCAGCAGATATTGGATGTCATGGGTGAGGCAGGCACCCGGCAAAGGACTGGAATGGATCGGCGAAATCAACCCAACCTCTAGCACCATTAACTTCACTCCTAGCCTGAAGGATAAGGTGTTCATCTCCAGGGACAACGCCAAGAATACACTGTACCTGCAGATGAGCAAGGTGAGATCTGAGGACACCGCCCTGTATTACTGTGCCAGAGGCAACTACTACAGGTATGGCGACGCCATGGACTACTGGGGCCAGGGCACATCAGTGACCGTGAGCTCC | EVKLLESGGGLVQPGGSLKLSCAASGFDFSRYWMSWVRQAPGKGLEWIGEINPTSSTINFTPSLKDKVFISRDNAKNTLYLQMSKVRSEDTALYYCARGNYYRYGDAMDYWGQGTSVTVSS | E132-S252 |
| HisTags + GCN4 linker | N/A | CATCATCACCATCACCACGGAGGTTCCGGTATGAAGCAGCTGGAGGACAAGGTCGAGGAGCTGCTGAGCAAGAACTACCACCTGGAGAATGAGAAGGCCCGCCTGGAGAAGCTGGTAGGCGAGAGGGGCGGATCCCACCATCACCATCACCATGCTGGTGGG | HHHHHHGGSGMKQLEDKVEELLSKNYHLENEKARLEKLVGERGGSHHHHHHAGG | H253-G306 |
| TP1107 (nanobody) | α-IgG1 Fc | CAGGTGCAGCTGGTGGAGTCCGGGGGAGGACTGGTGCAGCCTGGCGGCAGCCTGCGGCTGAGCTGTGCCGCCTCAGGCTTCACCTTTTCCGACACTTGGATGAATTGGGTGAGACAGGCACCCGGCAAAGGCCTGTATTGGATCAGCGCCATCAACCCTGACGGCGGAAATACCGCCTACGCCGATTCCGTGAAGGGAAGATTCACTATTAGCAGGGACAATGCCAAGAACATGGTGTATCTGCAGATGGACAATCTGCGCCCCGAGGACACCGCTATGTACTACTGTGCCAAGGGATGGGTGAGACTGCCTGATCCAGACCTGGTGAGAGGCCAGGGCACCCAGGTGACCGTGTCAAGCTGATAGGGATCCTGTCGTCGGATCTCCCCCGACCTC | QVQLVESGGGLVQPGGSLRLSCAASGFTFSDTWMNWVRQAPGKGLYWISAINPDGGNTAYADSVKGRFTISRDNAKNMVYLQMDNLRPEDTAMYYCAKGWVRLPDPDLVRGQGTQVTVSS | Q306-S426 |
| **CaptureBody-2** | α-IgG1 Fc x α-CD45 | CATCCTTTTTCTAGTAGCAACTGCAACCGGTGTACATTCCCAGGTGCAGCTGGTGGAGTCCGGGGGAGGACTGGTGCAGCCTGGCGGCAGCCTGCGGCTGAGCTGTGCCGCCTCAGGCTTCACCTTTTCCGACACTTGGATGAATTGGGTGAGACAGGCACCCGGCAAAGGCCTGTATTGGATCAGCGCCATCAACCCTGACGGCGGAAATACCGCCTACGCCGATTCCGTGAAGGGAAGATTCACTATTAGCAGGGACAATGCCAAGAACATGGTGTATCTGCAGATGGACAATCTGCGCCCCGAGGACACCGCTATGTACTACTGTGCCAAGGGATGGGTGAGACTGCCTGATCCAGACCTGGTGAGAGGCCAGGGCACCCAGGTGACCGTGTCAAGCCATCATCACCATCACCACGGAGGTTCCGGTATGAAGCAGCTGGAGGACAAGGTCGAGGAGCTGCTGAGCAAGAACTACCACCTGGAGAATGAGAAGGCCCGCCTGGAGAAGCTGGTAGGCGAGAGGGGCGGATCCCACCATCACCATCACCATGCTGGTGGGCAGGTGCAGCTGCAAGAGTCTGGCGGCGGCCTGGTGCAGACAGGTGGGAGCCTGACCCTGAGCTGCGTGGCCTCCGGCGGGACATTCAGCTCTTACGCCATGGGCTGGTTCAGACAGGCCCCAGGGAAGGAGAGGGAGTTCGTGGCCGCCATCGGCGGCAGCGGCGATTCTACCTATTATGCCGACAGCGTGAAAGGGAGATTCACCATCTCCAGAGATAACGCCAAGAATTCCGTGTACCTGCAGATGAACTCCCTGAAACCAGAGGACACAGCCGTGTACTACTGTCAGGCCGACCCTACTATGTTTCACAAGCTGTACTACGGAATTAACCCTAATGAGTATGACTACTGGGGCCAGGGCACCCTGGTGACAGTGAGCAGCTGATAGGGATCCTGTCGTCGGATCTCCCCCGACCTC | QVQLVESGGGLVQPGGSLRLSCAASGFTFSDTWMNWVRQAPGKGLYWISAINPDGGNTAYADSVKGRFTISRDNAKNMVYLQMDNLRPEDTAMYYCAKGWVRLPDPDLVRGQGTQVTVSSHHHHHHGGSGMKQLEDKVEELLSKNYHLENEKARLEKLVGERGGSHHHHHHAGGQVQLQESGGGLVQTGGSLTLSCVASGGTFSSYAMGWFRQAPGKEREFVAAIGGSGDSTYYADSVKGRFTISRDNAKNSVYLQMNSLKPEDTAVYYCQADPTMFHKLYYGINPNEYDYWGQGTLVTVSS | Q1-S302 |
| Mouse Ig heavy chain leader sequence | N/A | CATCCTTTTTCTAGTAGCAACTGCAACCGGTGTACATTCC | N/A | N/A |
| TP1107 (nanobody) | α-IgG1 Fc | CAGGTGCAGCTGGTGGAGTCCGGGGGAGGACTGGTGCAGCCTGGCGGCAGCCTGCGGCTGAGCTGTGCCGCCTCAGGCTTCACCTTTTCCGACACTTGGATGAATTGGGTGAGACAGGCACCCGGCAAAGGCCTGTATTGGATCAGCGCCATCAACCCTGACGGCGGAAATACCGCCTACGCCGATTCCGTGAAGGGAAGATTCACTATTAGCAGGGACAATGCCAAGAACATGGTGTATCTGCAGATGGACAATCTGCGCCCCGAGGACACCGCTATGTACTACTGTGCCAAGGGATGGGTGAGACTGCCTGATCCAGACCTGGTGAGAGGCCAGGGCACCCAGGTGACCGTGTCAAGC | QVQLVESGGGLVQPGGSLRLSCAASGFTFSDTWMNWVRQAPGKGLYWISAINPDGGNTAYADSVKGRFTISRDNAKNMVYLQMDNLRPEDTAMYYCAKGWVRLPDPDLVRGQGTQVTVSS | Q1-S120 |
| HisTags + GCN4 linker | N/A | CATCATCACCATCACCACGGAGGTTCCGGTATGAAGCAGCTGGAGGACAAGGTCGAGGAGCTGCTGAGCAAGAACTACCACCTGGAGAATGAGAAGGCCCGCCTGGAGAAGCTGGTAGGCGAGAGGGGCGGATCCCACCATCACCATCACCATGCTGGTGGG | HHHHHHGGSGMKQLEDKVEELLSKNYHLENEKARLEKLVGERGGSHHHHHHAGG | H121-G174 |
| 2H5 (nanobody) | α-CD45 | CAGGTGCAGCTGCAAGAGTCTGGCGGCGGCCTGGTGCAGACAGGTGGGAGCCTGACCCTGAGCTGCGTGGCCTCCGGCGGGACATTCAGCTCTTACGCCATGGGCTGGTTCAGACAGGCCCCAGGGAAGGAGAGGGAGTTCGTGGCCGCCATCGGCGGCAGCGGCGATTCTACCTATTATGCCGACAGCGTGAAAGGGAGATTCACCATCTCCAGAGATAACGCCAAGAATTCCGTGTACCTGCAGATGAACTCCCTGAAACCAGAGGACACAGCCGTGTACTACTGTCAGGCCGACCCTACTATGTTTCACAAGCTGTACTACGGAATTAACCCTAATGAGTATGACTACTGGGGCCAGGGCACCCTGGTGACAGTGAGCAGCTGATAGGGATCCTGTCGTCGGATCTCCCCCGACCTC | QVQLQESGGGLVQTGGSLTLSCVASGGTFSSYAMGWFRQAPGKEREFVAAIGGSGDSTYYADSVKGRFTISRDNAKNSVYLQMNSLKPEDTAVYYCQADPTMFHKLYYGINPNEYDYWGQGTLVTVSS | Q175-S302 |
| **CaptureBody-3** | α-IgG(κ) x α-CD45 | CATCCTTTTTCTAGTAGCAACTGCAACCGGTGTACATTCCCAGGTGCAGCTGGTGGAGAGCGGGGGCGGCTGGGTGCAGCCCGGAGGCAGTCTGAGGCTGAGCTGCGCCGCCAGCGGGTTTACCTTCTCCGACACAGCCATGATGTGGGTGAGACAGGCCCCTGGCAAGGGCCGCGAGTGGGTGGCCGCCATTGACACTGGCGGAGGTTACACCTACTACGCCGATTCCGTGAAGGGAAGGTTTACTATCTCCAGAGACAACGCCAAAAACACCCTGTACCTGCAGATGAACTCCCTGAAACCCGAGGACACCGCCAGATACTACTGTGCCAAGACTTACTCTGGAAATTATTACAGCAACTACACCGTGGCAAATTACGGCACCACCGGCCGGGGCACCCTGGTGACAGTGTCCTCCCATCATCACCATCACCACGGAGGTTCCGGTATGAAGCAGCTGGAGGACAAGGTCGAGGAGCTGCTGAGCAAGAACTACCACCTGGAGAATGAGAAGGCCCGCCTGGAGAAGCTGGTAGGCGAGAGGGGCGGATCCCACCATCACCATCACCATGCTGGTGGGCAGGTGCAGCTGCAAGAGTCTGGCGGCGGCCTGGTGCAGACAGGTGGGAGCCTGACCCTGAGCTGCGTGGCCTCCGGCGGGACATTCAGCTCTTACGCCATGGGCTGGTTCAGACAGGCCCCAGGGAAGGAGAGGGAGTTCGTGGCCGCCATCGGCGGCAGCGGCGATTCTACCTATTATGCCGACAGCGTGAAAGGGAGATTCACCATCTCCAGAGATAACGCCAAGAATTCCGTGTACCTGCAGATGAACTCCCTGAAACCAGAGGACACAGCCGTGTACTACTGTCAGGCCGACCCTACTATGTTTCACAAGCTGTACTACGGAATTAACCCTAATGAGTATGACTACTGGGGCCAGGGCACCCTGGTGACAGTGAGCAGCTGATAGGGATCCTGTCGTCGGATCTCCCCCGACCTC | QVQLVESGGGWVQPGGSLRLSCAASGFTFSDTAMMWVRQAPGKGREWVAAIDTGGGYTYYADSVKGRFTISRDNAKNTLYLQMNSLKPEDTARYYCAKTYSGNYYSNYTVANYGTTGRGTLVTVSSHHHHHHGGSGMKQLEDKVEELLSKNYHLENEKARLEKLVGERGGSHHHHHHAGGQVQLQESGGGLVQTGGSLTLSCVASGGTFSSYAMGWFRQAPGKEREFVAAIGGSGDSTYYADSVKGRFTISRDNAKNSVYLQMNSLKPEDTAVYYCQADPTMFHKLYYGINPNEYDYWGQGTLVTVSS | Q1-S308 |
| Mouse Ig heavy chain leader sequence | N/A | CATCCTTTTTCTAGTAGCAACTGCAACCGGTGTACATTCC | N/A | N/A |
| TP1170 (nanobody) | α-IgG(κ) | CAGGTGCAGCTGGTGGAGAGCGGGGGCGGCTGGGTGCAGCCCGGAGGCAGTCTGAGGCTGAGCTGCGCCGCCAGCGGGTTTACCTTCTCCGACACAGCCATGATGTGGGTGAGACAGGCCCCTGGCAAGGGCCGCGAGTGGGTGGCCGCCATTGACACTGGCGGAGGTTACACCTACTACGCCGATTCCGTGAAGGGAAGGTTTACTATCTCCAGAGACAACGCCAAAAACACCCTGTACCTGCAGATGAACTCCCTGAAACCCGAGGACACCGCCAGATACTACTGTGCCAAGACTTACTCTGGAAATTATTACAGCAACTACACCGTGGCAAATTACGGCACCACCGGCCGGGGCACCCTGGTGACAGTGTCCTCC | QVQLVESGGGWVQPGGSLRLSCAASGFTFSDTAMMWVRQAPGKGREWVAAIDTGGGYTYYADSVKGRFTISRDNAKNTLYLQMNSLKPEDTARYYCAKTYSGNYYSNYTVANYGTTGRGTLVTVSS | Q1-S126 |
| HisTags + GCN4 linker | N/A | CATCATCACCATCACCACGGAGGTTCCGGTATGAAGCAGCTGGAGGACAAGGTCGAGGAGCTGCTGAGCAAGAACTACCACCTGGAGAATGAGAAGGCCCGCCTGGAGAAGCTGGTAGGCGAGAGGGGCGGATCCCACCATCACCATCACCATGCTGGTGGG | HHHHHHGGSGMKQLEDKVEELLSKNYHLENEKARLEKLVGERGGSHHHHHHAGG | H127-G180 |
| 2H5 (nanobody) | α-CD45 | CAGGTGCAGCTGCAAGAGTCTGGCGGCGGCCTGGTGCAGACAGGTGGGAGCCTGACCCTGAGCTGCGTGGCCTCCGGCGGGACATTCAGCTCTTACGCCATGGGCTGGTTCAGACAGGCCCCAGGGAAGGAGAGGGAGTTCGTGGCCGCCATCGGCGGCAGCGGCGATTCTACCTATTATGCCGACAGCGTGAAAGGGAGATTCACCATCTCCAGAGATAACGCCAAGAATTCCGTGTACCTGCAGATGAACTCCCTGAAACCAGAGGACACAGCCGTGTACTACTGTCAGGCCGACCCTACTATGTTTCACAAGCTGTACTACGGAATTAACCCTAATGAGTATGACTACTGGGGCCAGGGCACCCTGGTGACAGTGAGCAGCTGATAGGGATCCTGTCGTCGGATCTCCCCCGACCTC | QVQLQESGGGLVQTGGSLTLSCVASGGTFSSYAMGWFRQAPGKEREFVAAIGGSGDSTYYADSVKGRFTISRDNAKNSVYLQMNSLKPEDTAVYYCQADPTMFHKLYYGINPNEYDYWGQGTLVTVSS | Q181-S308 |
| Original GCN4 Linker | N/A | ATGAAGCAGCTGGAGGACAAGATCGAGGAGCTGCTGAGCAAGATCTACCACCTGGAGAATGAAATCGCCCGCCTGAAGAAGCTGATTGGCGAGAGGGGC | MKQLEDKIEELLSKIYHLENEIARLKKLIGERG | M1-G33 |
| Post-Optimization GCN4 Linker | N/A | ATGAAGCAGCTGGAGGACAAGGTCGAGGAGCTGCTGAGCAAGAACTACCACCTGGAGAATGAGAAGGCCCGCCTGGAGAAGCTGGTAGGCGAGAGGGGC | MKQLEDKVEELLSKNYHLENEKARLEKLVGERG | M1-G33 |

**Supplementary Table 1: CaptureBody sequences, domain annotations, and amino acid modifications during GCN4 linker optimization**

| File | Donor | Condition | $DATE | Lymphocytes/PE His+ \| Median (YG1 (575)-A :: YG1 (575)-A) |
| --- | --- | --- | --- | --- |
| A01_BP0823685_No_Stain.fcs | BP0823685 | No Stain | 14-Oct-25 | 231 |
| A02_BP123190_No_Stain.fcs | BP123190 | No Stain | 14-Oct-25 | 271 |
| A03_BP0423145_No_Stain.fcs | BP0423145 | No Stain | 14-Oct-25 | 1179 |
| B01_BP0823685_Com_CD45_PE.fcs | BP0823685 | Commercial CD45 | 14-Oct-25 | 2.88E+06 |
| B02_BP123190_Com_CD45_PE.fcs | BP123190 | Commercial CD45 | 14-Oct-25 | 3.28E+06 |
| B03_BP0423145_Com_CD45_PE.fcs | BP0423145 | Commercial CD45 | 14-Oct-25 | 2.77E+06 |
| C01_BP0823685_Isotype_Only.fcs | BP0823685 | Isotype | 14-Oct-25 | 341 |
| C02_BP123190_Isotype_Only.fcs | BP123190 | Isotype | 14-Oct-25 | 371 |
| C03_BP0423145_Isotype_Only.fcs | BP0423145 | Isotype | 14-Oct-25 | 1189 |
| D01_BP0823685_muFc_scFV_nano_GCN4.fcs | BP0823685 | muFc scFv nano GCN4 | 14-Oct-25 | 2.20E+06 |
| D02_BP123190_muFc_scFV_nano_GCN4.fcs | BP123190 | muFc scFv nano GCN4 | 14-Oct-25 | 2.65E+06 |
| D03_BP0423145_muFc_scFV_nano_GCN4.fcs | BP0423145 | muFc scFv nano GCN4 | 14-Oct-25 | 1.44E+06 |
| E01_BP0823685_muFC_nano_nano_GCN4.fcs | BP0823685 | muFc nano nano GCN4 | 14-Oct-25 | 2.30E+06 |
| E02_BP123190_muFC_nano_nano_GCN4.fcs | BP123190 | muFc nano nano GCN4 | 14-Oct-25 | 2.90E+06 |
| E03_BP0423145_muFC_nano_nano_GCN4.fcs | BP0423145 | muFc nano nano GCN4 | 14-Oct-25 | 183590 |
| F01_BP0823685_muIgkappa_nano_nano_GCN4.fcs | BP0823685 | muIg kappa nano nano GCN4 | 14-Oct-25 | 2.11E+06 |
| F02_BP123190_muIgkappa_nano_nano_GCN4.fcs | BP123190 | muIg kappa nano nano GCN4 | 14-Oct-25 | 2.79E+06 |
| F03_BP0423145_muIgkappa_nano_nano_GCN4.fcs | BP0423145 | muIg kappa nano nano GCN4 | 14-Oct-25 | 9.22E+05 |
| B02_BP423145_No_Stain.fcs | BP0423145 | No Stain | 24-Oct-25 | 140 |
| C02_BP0423145_com_CD45.fcs | BP0423145 | Commercial CD45 | 24-Oct-25 | 2.90E+06 |
| D02_0423145_Isotype.fcs | BP0423145 | Isotype | 24-Oct-25 | 201 |
| E02_0423145_Isotype_Dimer1.fcs | BP0423145 | muFc scFv nano GCN4 | 24-Oct-25 | 1.74E+06 |
| F02_0423145_Isotype_Dimer4.fcs | BP0423145 | muFc nano nano GCN4 | 24-Oct-25 | 2.26E+06 |
| G02_0423145_Isotype_Dimer5.fcs | BP0423145 | muIg kappa nano nano GCN4 | 24-Oct-25 | 2.29E+06 |

| CaptureBody MFI Retention Per Donor | |
| --- | --- |
| **BP0823685** | |
| CB-1 | 76.4 |
| CB-2 | 79.8 |
| CB-3 | 73.3 |

| **BP123190** | |
| --- | --- |
| CB-1 | 80.8 |
| CB-2 | 88.4 |
| CB-3 | 85.0 |

| **BP0423145** | |
| --- | --- |
| CB-1 | 60.0 |
| CB-2 | 77.9 |
| CB-3 | 79.0 |

| CaptureBody Average MFI Retention | |
| --- | --- |
| CB-1 | 72.4 |
| CB-2 | 82.1 |
| CB-3 | 79.1 |

| In graph: | |
| --- | --- |
| dark | BP0823685 |
| medium | BP123190 |
| light | BP0423145 |
| red - not used in final analysis | |

**Supplementary Table 2: MFI reproduction of CaptureBodies and PE isotype compared to commercial anti-CD45 PE across 3 PBMC donors**

| **Fluorochrome** | **Group** | **File Name** | **Ungated / Size / Positive Median of UV16 (675)-A :: UV16 (675)-A** |
|---|---|---|---|
| **BUV661** | CB-1 | BUV661 A01.fcs | 241789 |
| **BUV661** | CB-2 | BUV661 A02.fcs | 608928 |
| **BUV661** | CB-3 | BUV661 A03.fcs | 527862 |
| **BUV661** | PBMC | BUV661 A05.fcs | 528613 |
| **BUV661** | Particles | BUV661 A06.fcs | 530400 |
| **Fluorochrome** | Group | File Name | Ungated / Size / Positive Median of UV18 (725)-A :: UV18 (725)-A |
| **BUV737** | CB-1 | BUV737 CD4 Depleted Dimer1.fcs | 142864 |
| **BUV737** | CB-2 | BUV737 CD4 Depleted Dimer4.fcs | 280024 |
| **BUV737** | CB-3 | BUV737 CD4 Depleted Dimer5.fcs | 205964 |
| **BUV737** | PBMC | BUV737 PBMC.fcs | 221813 |
| **BUV737** | Particles | BUV737 Beads.fcs | 228927 |
| **Fluorochrome** | Group | File Name | Ungated / Size / Positive Median of V16 (725)-A :: V16 (725)-A |
| **BV711** | CB-1 | BV711 B01.fcs | 148980 |
| **BV711** | CB-2 | BV711 B02.fcs | 301738 |
| **BV711** | CB-3 | BV711 B03.fcs | 272873 |
| **BV711** | PBMC | BV711 B05.fcs | 266568 |
| **BV711** | Particles | BV711 B06.fcs | 329411 |
| **Fluorochrome** | Group | File Name | Ungated / Size / Positive Median of B11 (700)-A :: B11 (700)-A |
| **PerCP-Cy5.5** | CB-1 | PerCP-Cy5.5 C01.fcs | 34985 |
| **PerCP-Cy5.5** | CB-2 | PerCP-Cy5.5 C02.fcs | 86065 |
| **PerCP-Cy5.5** | CB-3 | PerCP-Cy5.5 C03.fcs | 90463 |
| **PerCP-Cy5.5** | PBMC | PerCP-Cy5.5 C05.fcs | 54189 |
| **PerCP-Cy5.5** | Particles | PerCP-Cy5.5 C06.fcs | 79269 |
| **Fluorochrome** | Group | File Name | Ungated / Size / Positive Median of YG6 (675)-A :: YG6 (675)-A |
| **PE-Cy5** | CB-1 | PE-Cy5 D01.fcs | 1139216 |
| **PE-Cy5** | CB-2 | PE-Cy5 D02.fcs | 2358606 |
| **PE-Cy5** | CB-3 | PE-Cy5 D03.fcs | 2413527 |
| **PE-Cy5** | PBMC | PE-Cy5 D05.fcs | 1784688 |
| **PE-Cy5** | Particles | PE-Cy5 D06.fcs | 2534168 |

**Supplementary Table 3: MFI of positive populations of CaptureBodies with CD4 depleted PBMCs, unfractionated PBMCs, and compensation/unmixing particles with anti-CD4 fluorochromes (BUV661, BUV737, BV711, PerCP-Cy5.5, and PE-Cy5)**
